# Unveiling the hidden interactome of CRBN molecular glues with chemoproteomics

**DOI:** 10.1101/2024.09.11.612438

**Authors:** Kheewoong Baek, Rebecca J. Metivier, Shourya S. Roy Burman, Jonathan W. Bushman, Hojong Yoon, Ryan J. Lumpkin, Dinah M. Abeja, Megha Lakshminarayan, Hong Yue, Samuel Ojeda, Alyssa L. Verano, Nathanael S. Gray, Katherine A. Donovan, Eric S. Fischer

## Abstract

Targeted protein degradation and induced proximity refer to strategies that leverage the recruitment of proteins to facilitate their modification, regulation or degradation. As prospective design of glues remains challenging, unbiased discovery methods are needed to unveil hidden chemical targets. Here we establish a high throughput affinity purification mass spectrometry workflow in cell lysates for the unbiased identification of molecular glue targets. By mapping the targets of 20 CRBN-binding molecular glues, we identify 298 protein targets and demonstrate the utility of enrichment methods for identifying novel targets overlooked using established methods. We use a computational workflow to estimate target confidence and perform a biochemical screen to identify a lead compound for the new non-ZF target PPIL4. Our study provides a comprehensive inventory of targets chemically recruited to CRBN and delivers a robust and scalable workflow for identifying new drug-induced protein interactions in cell lysates.

## INTRODUCTION

Targeted Protein Degradation (TPD) represents a promising therapeutic approach to remove disease-associated proteins from cells^1,2^. The process of TPD involves hijacking the ubiquitylation machinery for the covalent attachment of ubiquitin molecules to a desired protein of interest, which in turn leads to degradation by the proteasome^3,4^. Ubiquitin-mediated TPD utilizes two types of small molecules, molecular glues^5^ and heterobifunctional degraders (also known as PROteolysis Targeting Chimeras, or PROTACs)^3^, both of which chemically induce ternary complex formation between a protein target and a ubiquitin E3 ligase, followed by proximity-driven ubiquitylation and subsequent degradation.

Despite the rapid growth of TPD as a therapeutic strategy, the discovery and development of effective degraders remains challenging. Heterobifunctional degraders rely on a linker to connect two binding warheads: one for the ligase and one for the protein of interest^3,6,7^. Although this modular design offers the flexibility to target any protein with a known binder, the resulting molecules often possess poor drug-like properties due to their large size. Molecular glues present a possible alternative due to their small size and improved drug-like properties. However, as they lack binding to their target protein and instead enhance protein-protein interactions (PPI) between a ligase and substrate, rational design of molecular glues is far more challenging^8^. Over the last decade, the discovery of new molecular glue degraders has largely relied on serendipity through phenotypic screening of large libraries of molecules and retrospective identification of their degradation targets. Although FDA-approved molecular glue degraders exist, including the immunomodulatory drugs (IMiDs) thalidomide, lenalidomide, and pomalidomide, they have all been characterized as molecular glues in retrospect after their serendipitous discovery. IMiD molecules bind to the CUL4-RBX1-DDB1-CRBN (CRL4^CRBN^) E3 ligase^5,9–13^ creating a favorable surface for new proteins (neo-substrates) to bind for induced degradation. Since this discovery, significant efforts into the design and screening of new IMiD analogs have revealed up to 50 neo-substrates in the public domain, all carrying a glycine containing β-hairpin structural degron^5,11,14–22^. Remarkably, computational modeling of the AlphaFold2 (AF2) structures available in the Protein Data Bank (PDB) suggest that we are just scraping the surface of what is targetable by these molecules^14^.

Given the mechanism of action (MoA) of degraders, global chemoproteomics has proven to be an effective tool for the identification of protein degradation targets^19,23–25^. Using this approach, the target space of degraders for multiple therapeutic target classes have been extensively mapped for tractable targets, including kinases^23,26,27^, bromo-domains^28–30^, HDACs^31^ and zinc finger (ZF) transcription factors^19^. Although this method has greatly expanded the repertoire of known targets, limited sensitivity has restricted the ability to identify proteins with low expression levels without screening libraries of cell lines or using target enrichment methods. This approach also remains blind to a key aspect of these molecules: the identification of proteins that are recruited to the ligase but do not ultimately get degraded. Such “non-degrading glue” targets may be subject to poor lysine accessibility, lack of degradative ubiquitin chain formation^32^, high deubiquitinase activity, poor proteasome access, or other resistance mechanisms. However, these targets still represent important therapeutic targets if these factors can be overcome to convert silent molecular glues into molecular glue degraders or functional modulators of the target.

Methods to identify chemically induced protein-protein interactions include immunoprecipitation mass spectrometry (IP-MS)^33^ and proximity labeling approaches coupled to mass spectrometry^34,35^. IP-MS approaches have been employed for the identification of direct protein interactions, whereas proximity labeling approaches are commonly employed for the mapping of proximity interactomes in cells and in vivo^34,36–38^, enabling the identification of protein interactions within a 10-30 nm radius of the epitope tagged protein of interest^34,36,37,39^. Although these in-cell methods have demonstrated successful identification of chemically induced interactions, they often require extensive fine tuning of various factors including noise, sensitivity, variability and scalability.

In this study, we establish a simple, robust and sensitive workflow to facilitate high throughput discovery of degrader-induced protein-protein interactions and develop them into selective tools and therapeutic candidates. We use this method to build a comprehensive inventory of 298 distinct protein targets recruited to CRBN including many new zinc finger (ZF) transcription factors and proteins from new target classes, including RNA-recognition motif (RRM) domain proteins. We evaluate the binding potential of these targets through structural alignment with IMiD-bound CRBN and performed biochemical and structural validation studies on a series of non-ZF targets. We then screened a library of ∼6000 IMiD analogs against a novel non-ZF target, PPIL4, identifying a selective lead degrader molecule, thereby presenting a blueprint for the effective discovery of novel molecular glue degraders.

## RESULTS

### Unbiased identification of degrader-induced interactors in-lysate

To establish a workflow for the identification of chemically induced protein-protein interactions, we set out to simplify traditional IP-MS methods. We hypothesized that we could create a controlled environment with reduced biological variability and enhanced scalability by establishing a workflow in cell lysate using spiked in recombinant protein as the bait. Our workflow harnesses small molecule degrader-induced ternary complex formation in cell lysate using recombinant FLAG-tagged CRBN in complex with DDB1 excluding the BPB domain (ΔB), which prevents CUL4 interaction to inhibit ubiquitylation of the recruited target, with a small molecule degrader. After incubation, we enrich with a highly selective antibody for the FLAG epitope tag followed by label free quantitative proteomic assessment to identify interactors (**Figure 1A**). To benchmark and explore the viability of this approach for identification of protein-protein interactions, we selected two representative degrader molecules that have been thoroughly profiled in published reports – pomalidomide, a IMiD molecular glue^19,40^ and SB1-G-187, a kinase-targeted heterobifunctional degrader^26^ (**Figure 1B**). We profiled these two molecules across two different cell lines, MOLT4 and Kelly, selected for their orthogonal expression profiles and broad coverage of known CRBN neo-substrates including IKZF1/3 (MOLT4) and SALL4 (Kelly)^19^. The pomalidomide screen revealed 11 different enriched proteins across these two cell lines (9 in MOLT4 and 4 in Kelly cells), which revealed three novel targets, ASS1, ZBED3 and ZNF219 (**Figure 1C**, **Figure S1A-B, Table S1**). We then validated recruitment of these three novel targets to CRBN using dose response immunoblot or TR-FRET analysis (**Figure S1C-D**).

**Figure 1.**
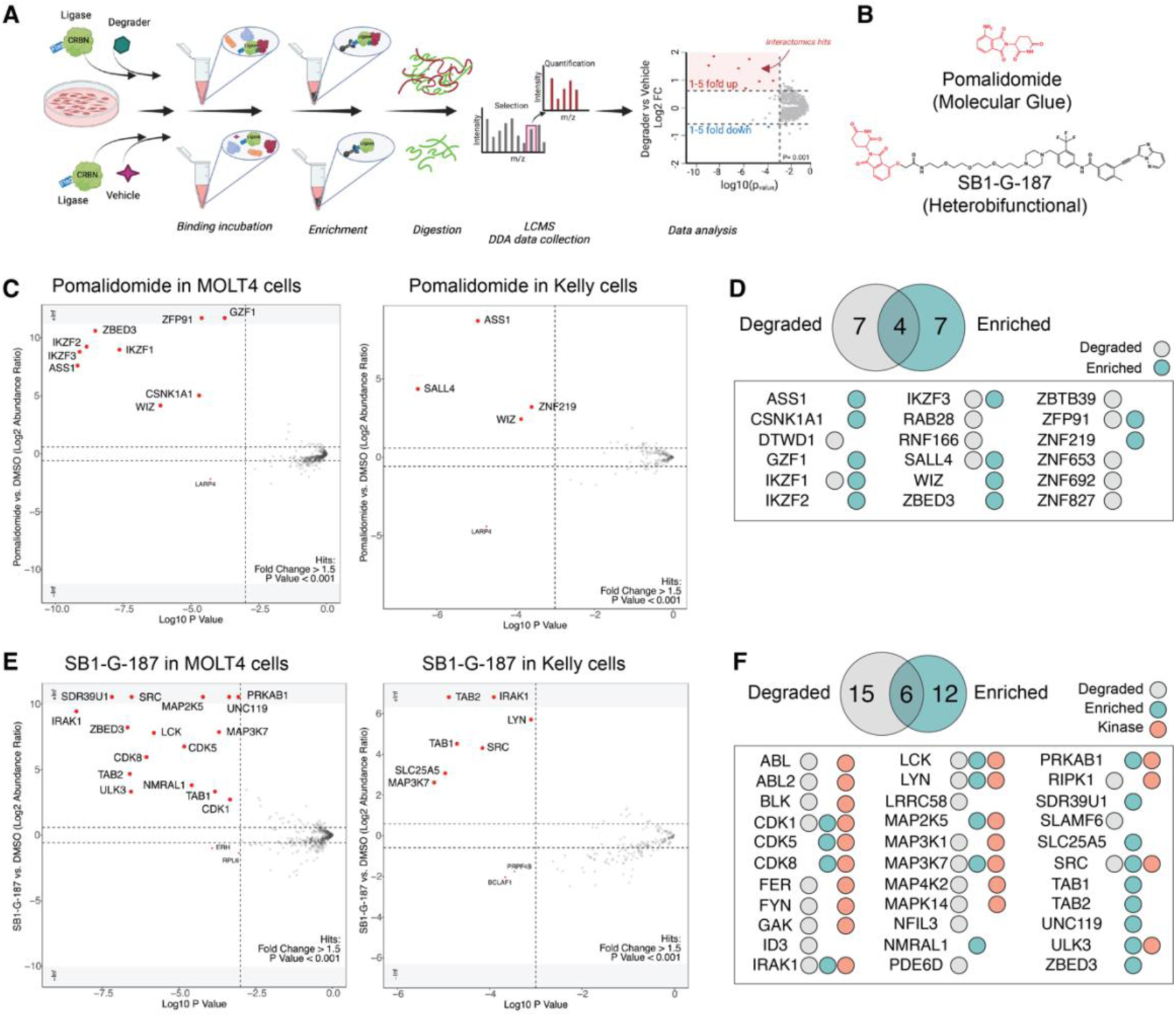
Proof of concept for target enrichment in-lysate. (**A**) Schematic representation of the first-generation enrichment-based quantitative proteomics workflow established for target enrichment and identification. (**B**) Chemical structures of degraders – Pomalidomide (molecular glue) and SB1-G-187 (heterobifunctional). (**C**) Scatterplots depicting relative protein abundance following Flag-CRBN-DDB1ΔB enrichment from in-lysate treatment with 1 µM Pomalidomide and recombinant Flag-CRBN-DDB1ΔB spike in. Left: MOLT4 cells and Right: Kelly Cells. Scatterplots display fold change in abundance to DMSO. Significant changes were assessed by moderated t-test as implemented in the limma package^80^ with log_2_ FC shown on the y-axis and negative log_10_ P-value on the x-axis. (**D**) Venn diagram showing unique and overlapping hits for Pomalidomide found in our enrichment study and in publicly available whole cell degradation data. (**E**) As in **C**, but with 1 µM SB1-G-187 treatment. (**F**) As in **D**, but with SB1-G-187 treatment.

To assess the overlap of these enriched targets with published degradation data for pomalidomide, we performed a hit comparison with publicly available global proteomics data (http://proteomics.fischerlab.org), which includes ten independent pomalidomide treatments spanning HEK293T, Kelly and MOLT4 cell lines (**Figure 1D**). Like our enrichment data, the global degradation data also maps 11 targets as degradable, however only 4 of these targets (IKZF1, IKZF3, ZFP91 and SALL4) overlap with those that we see enriched in this dataset. The SB1-G-187 kinase degrader screen identified 18 enriched targets across the two cell lines (16 in MOLT4 and 7 in Kelly cells) including multiple non-protein kinase targets which raised the question of how these proteins are being recruited to CRBN by a kinase degrader (**Figure 1E**, **Figure S1E-F, Table S1**). Assessment of the non-kinase targets revealed that several are known to form complexes with different kinases, such as TAB1 and TAB2 which form a functional kinase complex with MAP3K7^41,42^, and UNC119 which binds to myristoylated SRC to regulate cellular localization^43^. This data suggests that these non-kinases are being recruited to CRBN through piggybacking on their kinase binding partners. Of the other recruited non-kinase targets, ZBED3 is also identified in the pomalidomide treatment suggesting recruitment through the IMiD handle of the degrader and SDR39U1 was reported as a non-kinase target in a chemoproteomics profiling study of kinase inhibitor probes^44^. Next, we assessed the differences and overlap in hits between publicly available degradation data in MOLT4 and Kelly cells for kinase-targeted heterobifunctional degrader, SB1-G-187, and enrichment data (**Figure 1E**). We found 6 overlapping hits -all protein kinases - including CDK1, IRAK1, LCK, LYN, MAP3K7 and SRC. Like the pomalidomide data, we observe similar numbers of proteins identified in either degradation data (15) or enrichment data (12), demonstrating that these two methods complement each other to expand the target scope of these molecules. Together, the data collected for these two degrader molecules demonstrate the value of our workflow for identifying chemically induced protein-protein interactions invisible to degradation assays, while also highlighting opportunities for improving sensitivity.

### Mapping the interactomes of IMiD molecular glue degraders

Next, using the functional enrichment method as a basis, we set out to optimize and address the critical need for sensitivity and high throughput. IP-MS experiments typically require labor-intensive sample preparation steps which create a significant source of variability and lead to high background and false positive rates while also placing limits on the number of samples that can be prepared in parallel. To address these limitations, we automated the enrichment and sample preparation procedures to enable effective mapping of interaction targets across libraries of molecules at scale. We incorporated a cost effective Opentrons OT-2 liquid handling platform to automate the sample preparation process from addition of all immunoprecipitation components to tryptic digestion for 96 samples in parallel (**Figure 2A**). To address throughput and depth of the proteomics workflow, we took advantage of recent updates in instrumentation (timsTOF Pro2, Bruker) and acquisition methods (diaPASEF)^45^ that allow for significant improvements in sensitivity (**Figure 2A**). In contrast to the data-dependent acquisition (DDA) data collected in our proof-of-concept analysis (**Figure 1**), diaPASEF measures peptides by systematically sampling all precursor ions within a specified m/z range regardless of their abundance which enhances the reproducibility and depth of peptide coverage to allow for more accurate and robust quantification of peptides in complex samples. Comparison of the average numbers of proteins and peptides quantified between the DDA collection (**Figure 1A**) and diaPASEF collection (**Figure 2A**) revealed a >5-fold increase in proteins and a ∼9-fold increase in peptides quantified (**Figure S2A-B**) confirming a significant improvement in depth and sensitivity.

**Figure 2.**
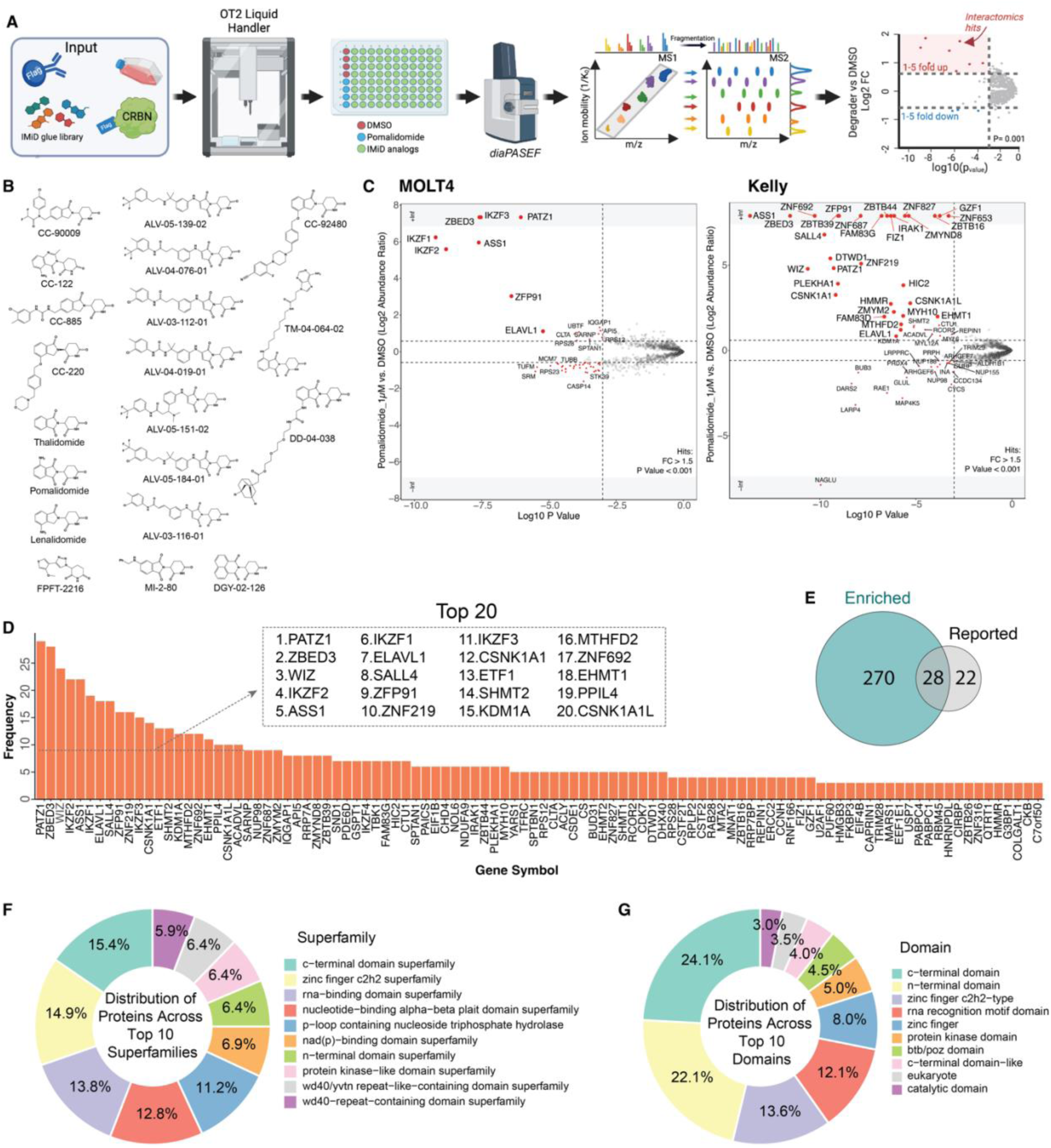
Unveiling and mapping CRBN recruited neo-substrates. (**A**) Schematic representation of the second-generation enrichment-based quantitative proteomics workflow established for target enrichment and identification. (**B**) Chemical structures of the 20 CRBN-based degraders profiled in this study. (**C**) Scatterplot depicting relative protein abundance following Flag-CRBN-DDB1ΔB enrichment from in-lysate treatment with degrader and recombinant Flag-CRBN-DDB1ΔB spike in. Scatterplot displays fold change in abundance to DMSO. Significant changes were assessed by moderated t-test as implemented in the limma package^80^ with log_2_ FC shown on the y-axis and negative log_10_ P-value on the x-axis. Scatterplots for all 21 treatments across MOLT4 and Kelly cells can be found in separate PDF’s “**Figures S3-4**”, representative example for a single treatment (Pomalidomide, 1 µM) is displayed here. (**D**) The number of independent IPs for which enrichment was observed for each target. Inset, the top 20 frequently enriched target proteins. (**E**) Venn diagram showing unique and overlapping hits found in our enrichment study and in published literature. (**F**) Donut chart representing the proportions of enriched proteins contained within the Top 10 different superfamily categories. (**G**) Donut chart representing the proportions of enriched proteins contained within the Top 10 different domain categories.

Work over the last several years has led to the identification of a growing list of ∼50 neo-substrates that are recruited to CRBN by IMiD analogs for chemically induced degradation^14^. Validated targets include a large number of C_2_H_2_ zinc finger (ZF) transcription factors such as IKZF1/3^5^, ZFP91^18^, or SALL4^15,19^, but only a few non-ZF proteins such as G1 to S phase transition protein 1 (GSPT1)^11^ and casein kinase 1 alpha (CK1α)^12,17^. These targets do not possess any similarity, but instead all share a common structural CRBN binding motif consisting of an 8-residue loop that connects the two strands of a β-hairpin and has a glycine at the sixth position (G-loop)^11,12,18^. Remarkably, a recently reported analysis of available AlphaFold2 (AF2) predicted structures for proteins in the human proteome uncovered over 2,500 proteins that harbor a G-loop potentially compatible with IMiD-recruitment to CRBN, with C_2_H_2_ ZF proteins revealing themselves as the most prevalent domain class, aligning with the dominance that this class has amongst the experimentally confirmed targets^14^. Due to the extensive range of proteins that are predicted to be chemically recruitable to CRBN, we asked how many of these proteins are already targeted by existing chemistry, but not yet identified due to lack of sensitivity of existing methods. To explore the range of proteins chemically recruited to CRBN, we screened a curated library of 20 different IMiD analogs through our automated lysate-based IP workflow. We assembled this library to incorporate a broad range of IMiD-based scaffolds including the parental FDA-approved IMiDs (thalidomide, lenalidomide, pomalidomide)^46,47^, where there is a high value to identifying new targets for drug repurposing efforts. We included a series of IMiD analogs that are undergoing clinical trials (CC-220, CC-92480, CC-90009)^48–50^ and molecules that have demonstrated promiscuity (FPFT-2216, CC-122)^51,52^. Finally, we included a series of in-house synthesized scaffolds developed in the context of targeting Helios (IKZF2)^53^ or part of an effort to diversify IMiDs with the addition of fragments on an extended linker (**Figure 2B**). We screened this library at 1 µM concentration across MOLT4 and Kelly cell lines (including a second 5 µM concentration for pomalidomide) and identified proteins that were enriched in the degrader compared to DMSO control IP treatment (**Figure 2C**, **Figures S3-4, Tables S2-3**). Using significance cutoffs of fold change (FC) >1.5, P-value <0.001 and combining the data from both cell lines, we identified a total of 298 enriched proteins (**Table S4**). We rationalized that the likelihood of observing the same proteins enriched as false positives across multiple treatments with similar IMiD analog molecules is low and therefore used ‘frequency of enrichment’ as a measure of confidence. We observed 102 proteins enriched in at least three independent IPs, and each of the top 5 proteins (PATZ1, ZBED3, WIZ, IKZF2 and ASS1) enriched in more than 20 independent IPs across the database (**Figure 2D**, **Table S4**). Surprisingly, although published reports have confirmed degradation of PATZ1, WIZ and IKZF2, none of these top 5 enriched proteins regularly feature amongst those proteins that we commonly see reported in existing unbiased screens of IMiD-based molecules indicating the orthogonal data generated by this profiling method, identifying targets that might otherwise be overlooked. ZBED3 and ASS1 showed frequent enrichment across our database without any prior reporting of degradation, even at concentrations up to 5 µM of pomalidomide (**Figure S2C, Table S4**), suggesting the first reported examples of targets that are chemically glued to CRBN but lack productive degradation, thus emphasizing the benefit of alternative binding focused approaches for target identification. Also, important to note is that the new targets identified in this study are not only targets of new IMiD analogs but are also identified as targets of IMiDs in clinical trials and with FDA approval.

To assess the fraction of newly identified IMiD targets, we compared the 298 enriched proteins to a list of literature reported targets and discovered an overlap of only 28 targets. We identified 270 novel targets and found only 22 targets were reported in the literature but not identified as hits in our study (**Figure 2E**, **Table S4**)^14^. Considering the prevalence of C_2_H_2_ ZF transcription factors amongst reported IMiD targets, we asked whether this dominance holds true across our extended list of targets. To assess this, we extracted superfamily, family and domain information from curated databases including InterPro^54,55^, Uniprot^56^ and Superfamily^57^ to categorize the targets based on studied features (**Figure 2F-G**, **Table S4**). Of the 298 targets identified, after C-terminal domain classification, the C_2_H_2_ ZF superfamily represents the largest segment, accounting for >14% of the targets in the top 10 enriched superfamilies. This is followed by RNA-recognition motif domain proteins (RRM, >13%) and nucleotide-binding alpha-beta plait domain superfamilies (α-β plait domain, >12%). Notably, protein kinase-like domain proteins also features on this top 10 list of superfamilies (kinase-like domain, >6.4%) which aligns with our knowledge that kinases can be targets of IMiD molecular glues (eg, CSNK1A1 or WEE1)^12,17,58^ and suggests that molecular glues may be a viable alternative to PROTACs, which are currently widely explored for kinase targeting. Exploration of the top 10 domain classifications across the dataset shows a similar trend with C_2_H_2_ ZF, RRM, ZF, protein kinases and BTB/POZ domains showing the highest representation across the targets identified (**Figure 2G**).

This dataset builds upon previous identifications of protein kinases as targets of IMiD molecules^17,22,59^, and further extends the kinase list adding CDK7, IRAK1 and TBK1 as novel putative molecular glue targets. It also broadens the scope of tractable targets by introducing multiple new families as targetable by CRBN-based molecular glues, illustrating the extensive potential of these molecules. Through the application of our unbiased target enrichment workflow, we have significantly increased the number of experimentally detected IMiD targets, expanding beyond the C_2_H_2_ ZF protein family to a wide range of protein families including protein kinases and proteins involved in RNA metabolism.

### Zinc finger transcription factors enriched among targets

To validate the 270 previously unreported targets, we sought to establish a computational screening pipeline to score the compatibility of targets for recruitment to CRBN. Structural studies on IMiD-mediated CRBN neo-substrates, both natural and designed, have established the common G-loop motif that is recognized by the CRBN-IMiD complex^12,60^. We used MASTER^61^ to mine the AF2 database^62^ for proteins containing G-loops with similar backbone architecture to the G-loop in known neo-substrate CSNK1A1 (PDB: 5FQD, aa35-42) resulting in a set of 46,040 loops across 10,926 proteins (**Figure 3A**).

**Figure 3.**
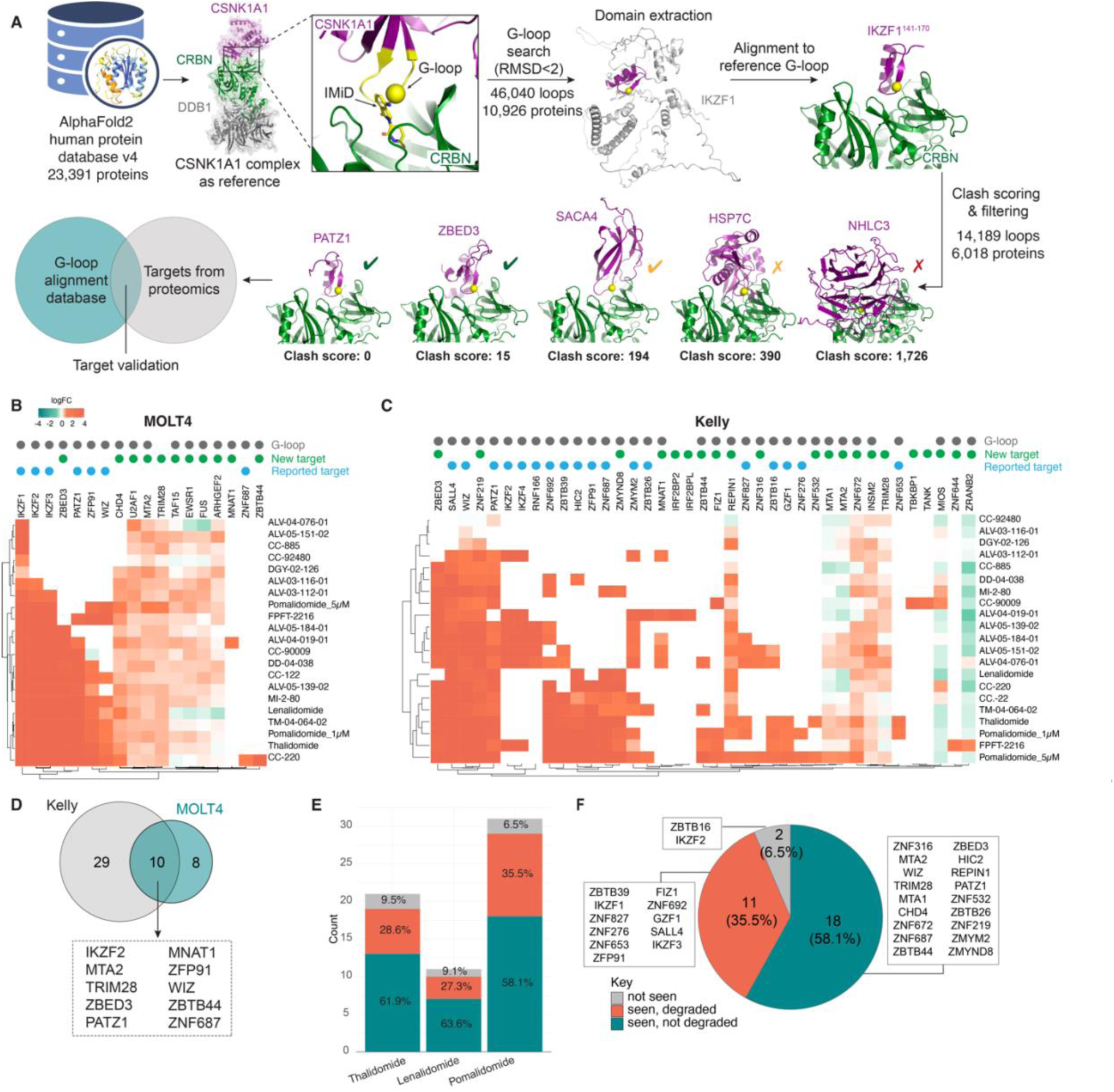
Structural alignment and assessment of ZF CRBN neo-substrates. (**A**) Schematic representation of the computational workflow established for AF2 G-loop binding compatibility with CRBN-IMiD. (**B**) Heatmap displaying the log2 fold change (log2 FC) of significant (P-value <0.001) molecular glue dependent ZF targets in MOLT4 cells. White space in the heatmap corresponds to log2FC = 0 or no quantification. Previously reported targets are marked with a blue dot, newly reported targets are marked with a green dot and targets with a structural G-loop are marked with a gray dot. Significant changes were assessed by moderated t-test as implemented in the limma package^80^. (**C**) As in **B**, but with Kelly cells. (**D**) Venn diagram showing unique and overlapping ZF hits comparing MOLT4 and Kelly cell targets. (**E**) Stacked bar plot showing the proportion of targets complexed and degraded by each of the indicated IMiD molecules. “not seen” indicates enriched targets were not quantified in global proteomics studies. “seen, degraded” indicates enriched targets quantified and reported as degraded in global proteomics. “seen, not degraded” indicates enriched targets were quantified but not degraded in global proteomics ^19^. (**F**) Pie chart displays the IMiD-grouped data from **E**.

Due to structural constraints, not all these proteins are compatible with CRBN. To identify CRBN-compatible proteins, we first extracted domains containing the G-loops based on domain definitions from DPAM, a tool that parses domains from AF models based upon predicted aligned errors (PAE) and evolutionary classification^63^. Next, we aligned the domains to our reference CSNK1A1-IMiD-CRBN-DDB1ΔB structure based on the G-loop and calculated a clash score. We used the van der Waals force term for interchain contacts in Rosetta’s low-resolution mode^64^ to obtain a side-chain independent clash estimate. Out of 16 known neo-substrates with validated G-loops (**Table S5, Figure S5**), 15 had clash scores below 2, while ZNF654^40^ had a score of 172, indicating a minor clash. The clash was caused by a low confidence region in the AF2 structure and could be resolved by relaxing the complex with Rosetta (**Figure S6A**)^65^. On the other hand, a protein with no evidence supporting it being a neo-substrate, PAAF1, had a major clash with a score of 1,551 which could not be resolved by relaxation (**Figure S6B**). Based on these examples, and analysis of the clash scores of all hit proteins containing a clear structural G-loop in AF2 (**Figure S6C)**, we filtered out domains with scores greater than 200 resulting in a list of 14,189 loops across 6,018 proteins with nonexistent, or marginal clashes with CRBN (**Figure 3A**, **Table S5**).

Of the 298 total enriched candidates,199 were found to have a clear structural G-loop with 162 having a clash score below 200. Given the high proportion of ZF proteins identified as targets across this enrichment database (**Figure 2F**), we mapped the fold change in enrichment for proteins with an annotated ZF domain across all 20 degraders for both MOLT4 (**Figure 3B**) and Kelly cells (**Figure 3C**). Across these two cell lines, we identified 19 previously reported and 28 new neo-substrates as chemically recruited to CRBN. We then used our G-loop database to inform on which of these targets have a tractable G-loop and found that only five of the 57 identified targets do not contain a structural G-loop (**Figure 3C**, **Table S5**). Given what we know about the recruitment and binding of CRBN neo-substrates, targets usually bind through a dominant structural hairpin. Since we do not have validated degron information for all these ZF targets, we assumed that the G-loop with the lowest clash score has the highest likelihood to bind and therefore proceeded with evaluation of a single G-loop for each target. To gauge how the clash scores for these ZF targets compare to all hits in the G-loop database, we compared the clash scores for our ZF targets to those of all hits (**Figure S6C**) demonstrating a pronounced trend towards lower scores for ZF targets suggesting fewer unfavorable interactions (**Figure S5A-B**). Notably, when we explored the ZF hits with higher clash scores (>10) and >3 hit frequency, we realized that almost all of these have a reported association with at least one of the validated hits – ZMYND8 (cs 455, binds to ZNF687), and RNF166 (cs 17, binds to ZNF653/ZBTB39/ZNF827) – which also offers the possibility that these proteins could be collateral targets, recruited via piggybacking on their binding partners, the direct binders (**Table S4**). Finally, we compared ZF targets across the two cell lines as an additional means for validation, and found 10 overlapping proteins, 6 of which are novel recruited targets (ZBED3, MNAT1, MTA2, ZBTB44, TRIM28) (**Figure 3D**).

There are many factors to take into consideration when looking to predict target degradability, such as ternary complex formation^26,31,66^ and target ubiquitylation^12,32,67–69^, and multiple studies have placed an emphasis on exploring their role in driving productive degradation^70,71^. For degrader-induced degradation to occur, a ternary complex consisting of ligase-degrader-target needs to form for proximity-mediated ubiquitin transfer to the target protein. Because ternary complex formation is necessary for successful protein degradation, we set out to explore the relationship between ternary complex formation and degradation for ZF targets identified in this study. We focused our evaluation on the parental IMiD molecules which have been subjected to degradation target profiling using unbiased global proteomics analysis across a panel of four cell lines (SK-N-DZ, Kelly, MM.1S, hES)^19^. Comparison of the enriched ZF targets to the published degradation data shows a consistent trend across the three IMiDs where only ∼30% of the enriched targets that were quantified in global proteomics studies were degraded, with ∼ 60% of the targets quantified but not reported as degraded (**Figure 3E**, **Table S5**). The data was then grouped to allow a global comparison of the enriched versus degraded IMiD targets. The comparison revealed that of the 31 ZF targets enriched across these three molecular glues, only 11 of the 29 proteins quantified in global proteomics experiments were found to be degraded (**Figure 3F**, **Table S5**). 18 proteins were quantified in global proteomics but were not identified as degraded. This prompted us to question whether these targets were resistant to degradation by IMiDs and their analogs, or if they were not identified as degraded due to experimental limitations such as inadequate sensitivity to detect minor changes in protein abundance, rapid protein turnover or suboptimal experimental conditions. We found that although several of the targets (WIZ, PATZ1, ZNF687, ZMYM2 and HIC2) were not reported as degraded in Donovan et al.^19^, they have since been reported as degraded in other published studies^40,72,73^ confirming that IPs provide a complementary approach able to overcome limitations in sensitivity. The absence of degradation data for the remaining targets could imply that these targets are resistant to degradation, or similar to the above proteins, the appropriate degradation experiment has yet to be performed. These data demonstrate that our IP workflow provides a significant advantage over global proteomics analysis by enabling selective isolation and enrichment of targets that may be below the change in abundance threshold for consistent identification with global proteomics approaches.

### IMiD derived molecular glues recruit hundreds of non-zinc finger proteins

The largest target class of CRBN neo-substrates today are ZF containing proteins, however, of the ∼20,000 proteins in the human proteome, ZF containing proteins only make up a relatively small proportion with about ∼1700 ZF proteins reported^74^. So far, only a handful of targets are reported to lack a ZF motif, which includes GSPT1^11^, CK1α^12,17^, PDE6D^75^, and RAB28^75^. With this in mind, we examined our list of targeted proteins with a focus on those that do not contain a reported ZF domain and found 251 non-ZF proteins enriched across the IP dataset (**Figure 4A**, **Table S4**). These non-ZF proteins include a wide range of families such as protein kinases (IRAK1, TBK1, CDK7), RNA recognition motif proteins (ELAVL1, PPIL4, CSTF2, RBM45), metabolic enzymes (ASS1, PAICS, ACLY, CS, ACADVL), translational proteins (MARS1, ETF1, EEF1E1, EIF4B) and more spanning different biological pathways. To assist in establishing confidence in some of these targets, we performed a comparison of the non-ZF targets enriched in the two tested cell lines and found 39 targets were identified in both MOLT4 and Kelly cells, including the four above mentioned targets (**Figure 4A-B**). We then assessed the AF2 structures of each of these 39 proteins and found that almost all of them (33/39) contain a structural G-loop (**Figure 4B**, **Figure S5, Table S5**).

**Figure 4.**
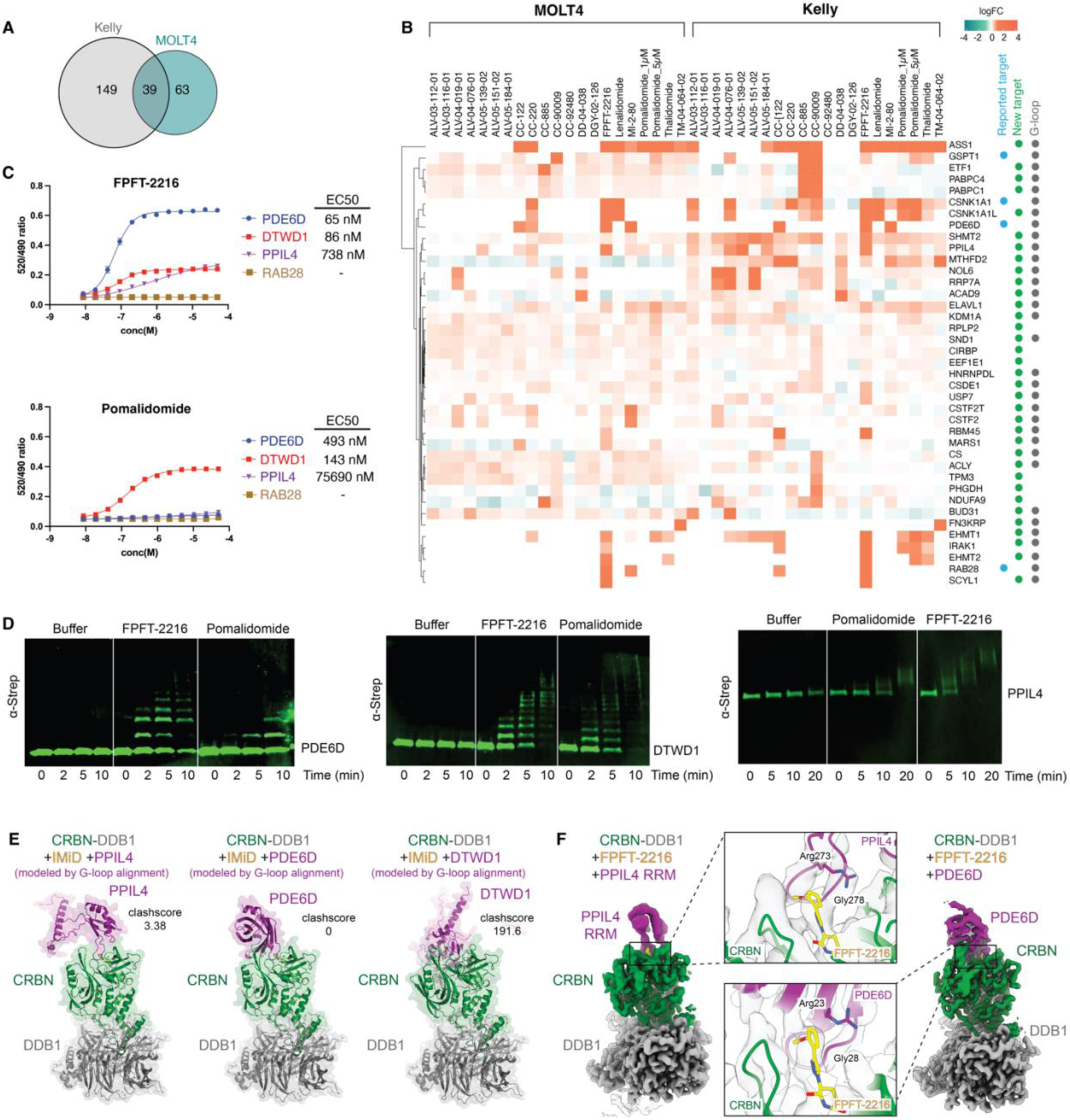
Assessment and validation of CRBN non-ZF neo-substrates. (**A**) Venn diagram showing unique and overlapping non-ZF hits comparing MOLT4 and Kelly cell targets. (**B**) Heatmap displaying the log2 fold change (log2 FC) for the 39 overlapping hits from **A**. White space in the heatmap corresponds to log2FC = 0 or no quantification. Previously reported targets are marked with a blue dot, newly reported targets are marked with a green dot and targets with a structural G-loop are marked with a gray dot. Significant changes were assessed by moderated t-test as implemented in the limma package^80^. (**C**) TR-FRET with titration of FPFT-2216 or pomalidomide to N-terminally biotinylated FL PDE6D, DTWD1, PPIL4 or RAB28 at 20 nM, incubated with terbium-streptavidin at 2 nM to monitor binding to GFP-CRBN-DDB1ΔB at 200 nM. Values were determined by technical replicates of n=2. **(D)** Immunoblots of ubiquitylation assay establishing PDE6D, DTWD1 and PPIL4 as FPFT-2216 and pomalidomide-induced neo-substrates of CRBN. (**E**) Structural G-loop alignment of AF2 PPIL4, PDE6D, and DTWD1 with CRBN-DDB1ΔB (PDB ID 5QFD, 6UML). Corresponding clash score is displayed. (**F**) Cryo-EM 3D reconstruction of PPIL4-RRM bound in ternary complex with FPFT-2216-CRBN-DDB1 FL, and PDE6D bound with FPFT-2216-CRBN-DDB1ΔB. Maps are postprocessed with DeepEMhancer^81^. Inset of each shows the potential binding mode of action of FPFT-2216 engaging PPIL4 or PDE6D via neo-substrate G-loop and its interacting residues.

Given the large number of non-ZF targets identified in this study and the lack of emphasis in the public domain with regards to non-ZF CRBN neo-substrates, we selected a series of non-ZF proteins for further experimental validation. Firstly, to demonstrate that these neo-substrates are directly recruited to CRBN, we examined ternary complex formation using recombinant purified proteins. Using two of the more promiscuous molecular glues, pomalidomide and FPFT-2216, we tested previous reported degradation targets PDE6D, RAB28 and DTWD1, along with a newly discovered target PPIL4. Indeed, PDE6D, DTWD1, and PPIL4 formed compound dependent ternary complex with CRBN at varying effective concentrations (**Figure 4C**). However, RAB28, which was previously reported to be degraded by IMiDs^19^ and FPFT-2216^75^, did not show any evidence for direct binding to CRBN using purified proteins. Since RAB28 has previously been reported as a CRBN neo-substrate and consistently scored across our enrichment study, we explored whether there was any evidence suggesting that RAB28 could be a collateral target. Exploration of protein-protein interaction databases including BioPlex^33^ and STRING-DB^76^ revealed that RAB28 is known to bind to two validated IMiD-CRBN neo-substrates PDE6D and ZNF653 (**Table S4**), suggesting that RAB28 is likely an indirectly recruited target. These data demonstrate that in addition to identifying direct binders, we can also identify indirect binding partners that may be simultaneously recruited together with direct binding neo-substrates. Given that targeted protein degradation requires not only recruitment to CRBN, but also CRBN mediated ubiquitin transfer for degradation, we also monitored whether the recruited proteins can be ubiquitylated by CRL4^CRBN^. In vitro ubiquitylation assays showed robust ubiquitin modification on all 3 recruited non-ZF proteins in the presence of pomalidomide or FPFT-2216 (**Figure 4D**). In addition, all three of these targets were degraded in response to IMiD treatment as observed by global proteomics analysis (**Figure S6D, Table S5**). Using structural G-loop alignments, we then assessed the potential for each of these three proteins to bind to IMiD-CRBN and found that all three proteins had a G-loop with a clash score of <200 (**Figure 4E**). However, the aligned clash score for DTWD1 was relatively high and approaching the upper 200 threshold (cs 198). We performed relaxation with Rosetta and found that this reduced the clash score to 1.58 by allowing minor shifts in the overall conformation while retaining the structural G-loop (**Figure S6E**). This process demonstrates that in some cases, clash scores can be relieved through minor structural rearrangements using Rosetta relax.

To expand our understanding of the recruitment of non-ZF targets, we determined cryo-EM structures of CRBN-DDB1ΔB-FPFT-2216 bound to PPIL4 and PDE6D, respectively (**Figure 4F**, **Figure S7**, **Table 1**). The complex structures were both refined to a global resolution of around 3.4 Å and the quality of the resulting maps were sufficient to dock the complex components, but the flexibly tethered PPIL4 resulted in a lower local resolution. We were able to observe PPIL4 engagement with FPFT-2216-CRBN via its Gly278 harboring G-loop as expected from the G-loop alignment, as well as for PDE6D via its Gly28 G-loop.

**Table 1.** Data collection and refinement statistics for cryo-EM datasets, related to Figures 4 and S7.

|  | CRBN-DDB1<br>FPFT-2216<br>PPIL4 RRM | CRBN-DDB1ΔB<br>FPFT-2216<br>PDE6D<br>Consensus refine | CRBN-DDB1ΔB<br>FPFT-2216<br>PDE6D<br>Local refine |
| --- | --- | --- | --- |
| Microscope | Talos Arctica | Talos Arctica |  |
| Voltage (kV) | 200 | 200 |  |
| Camera | Gatan K3 | Gatan K3 |  |
| Magnification (×) | 36,000 | 36,000 |  |
| Pixel size (Å) | 1.1 | 1.1 |  |
| Total electron exposure (e<sup>-</sup>/Å<sup>2</sup>) | 54 | 51.2 |  |
| Number of frames | 40 | 50 |  |
| Defocus range (μm) | −2.0 to −0.8 | −2.0 to −0.8 |  |
| Data collection software | SerialEM4.1b |  |  |
| Micrographs collected | 4,170 | 4,474 |  |
| Total extracted particles | 3,198,055 | 3,393,589 |  |
| EMDB accession code | EMD-XXXXXX | EMD-XXXXXX | EMD-XXXXXX |
| PDB accession code | PDB-XXXX | PDB-XXXX |  |
| Final particles used | 219,802 | 515,636 | 515,636 |
| Map resolution (Å, FSC 0.143) | 3.5 | 3.3 | 3.4 |
| FSC threshold | 0.143 | 0.143 | 0.143 |
| Model composition |  |  |  |
| Protein residues | 1211 | 1260 |  |
| Ligands | 2 | 2 |  |
| Refinement package | phenix.real_space_refine |  |  |
| Model-to-map CC | 0.72 | 0.78 |  |
| Model-to-map FSC (Å, FSC 0.5) | 3.8 | 3.5 |  |
| Mean *B* factors (Å<sup>2</sup>) |  |  |  |
| Protein | 63.69 | 98.29 |  |
| Ligand | 69.37 | 118.85 |  |
| Water | — | — |  |
| Bond root-mean-square deviation (RMSD) |  |  |  |
| Lengths (Å) | 0.005 | 0.005 |  |

|  | **CRBN-DDB1**<br>**FPFT-2216**<br>**PPIL4 RRM** | **CRBN-DDB1ΔB**<br>**FPFT-2216**<br>**PDE6D**<br>**Consensus refine** | **CRBN-DDB1ΔB**<br>**FPFT-2216**<br>**PDE6D**<br>**Local refine** |
| --- | --- | --- | --- |
| Angles (°) | 0.615 | 0.797 |  |
| **Validation** |  |  |  |
| MolProbity score | 2.05 | 1.53 |  |
| Clash score | 11.78 | 6.37 |  |
| Rotamer outliers (%) | 1.27 | 0.11 |  |
| CaBLAM outliers (%) | 2.76 | 1.54 |  |
| **Ramachandran plot** |  |  |  |
| Favored (%) | 94.26 | 96.96 |  |
| Allowed (%) | 5.74 | 3.04 |  |
| Disallowed (%) | 0.00 | 0.00 |  |

Furthermore, overall density allowed fitting of FPFT-2216 in bulk although the reduced resolution in that region did not permit exact positioning of the molecule. Nevertheless, we were able to see that the glutarimide ring engages CRBN’s binding pocket in a similar manner to other IMiD molecular glues. The triazole interacts with the backbone of the G-loop, and the methoxythiophene moiety potentially contacts both the PPIL4 backbone of the G-loop and Arg273. This suggests that the triazole and the methoxythiophene moieties could provide specificity elements to FPFT-2216 mediated neo-substrate recruitment. The methoxythiophene moiety also engaged Arg23 of PDE6D, indicating that FPFT-2216 might derive specificity in engaging an arginine residue from its neo-substrates. Analysis of the non-ZF targets of FPFT-2216 revealed several other proteins harboring an arginine or a lysine residue at this sequence location (PDE6D, SCYL1, RBM45, PPIL4). Finally, we compared the experimental structure to the AF2 predicted G-loop aligned structure of PPIL4 (**Figure S6F**). The G-loop aligned structure of PPIL4 presented a clash score of 3.38, which showed the C-terminal region of CRBN around residue Arg373 to be clashing with PPIL4’s loop harboring residue Val250. Although the low resolution permitted only backbone level fitting of PPIL4, we observed that the cryo-EM structure revealed a minor shift in the RRM domain of PPIL4 to accommodate this minor clash suggested in the G-loop aligned structure while retaining overall conformational similarity of the G-loop (**Figure S6F**). Meanwhile, PDE6D retained overall similar conformation with minor shifts that did not alter the interaction with CRBN (**Figure S6G**).

These data demonstrate that RRM domain containing proteins represent a new class of proteins targetable through CRBN dependent molecular glues. Using structural modeling we increase confidence in these new targets while also providing a reminder that structural analysis and AF2 predicted structures are static models and although they provide excellent structural guidance, we need to keep in mind that proteins in solution are flexible and dynamic.

### Discovery of new and selective molecular glue for PPIL4

While the proteomics-based screening workflow identifies novel putative CRBN targets and provides initial chemical matter, it does not necessarily provide the best starting point for developing a chemical probe or therapeutic due to the limited number of molecules screened. We hypothesized that this limitation could be overcome by following up proteomics screening with a target centric screen of a larger CRBN binder library to identify the optimal chemical starting point. To test this, we set out to identify PPIL4 targeting molecular glues with improved selectivity and lacking the triazole moiety. We employed an IMiD molecular glue library consisting of ∼6000 compounds of various IMiD analogs that were either synthesized in-house or purchased externally. We screened this library against PPIL4 using TR-FRET to measure compound-induced PPIL4 recruitment to CRBN (**Figure 5A**). TR-FRET ratios were obtained by incubating the library with GFP fused CRBN-DDB1ΔB, biotinylated PPIL4, and Tb-labeled streptavidin that binds to the biotinylated PPIL4. The library was compared relative to the positive control, whereby the 520/490 ratio of FPFT-2216 at 10 µM was normalized as 1, and compounds were tested at 1.66 µM or 3.33 µM to find hits with equal or improved efficacy in directly recruiting PPIL4 to CRBN-DDB1ΔB. We were able to narrow down the library to two molecules that performed similar or better than FPFT-2216 (**Figure 5B)**. These lead compounds were subject to a full titration to assess recruitment efficacy by TR-FRET. Ultimately, after recognizing one of the two hits was due to autofluorescence, we were able to identify a molecule, Z6466608628, that produced a higher 520/490 ratio, and a better EC50 of 0.34 µM compared to FPFT-2216, measured at 1.05 µM in this experiment (**Figure 5C-D**).

**Figure 5.**
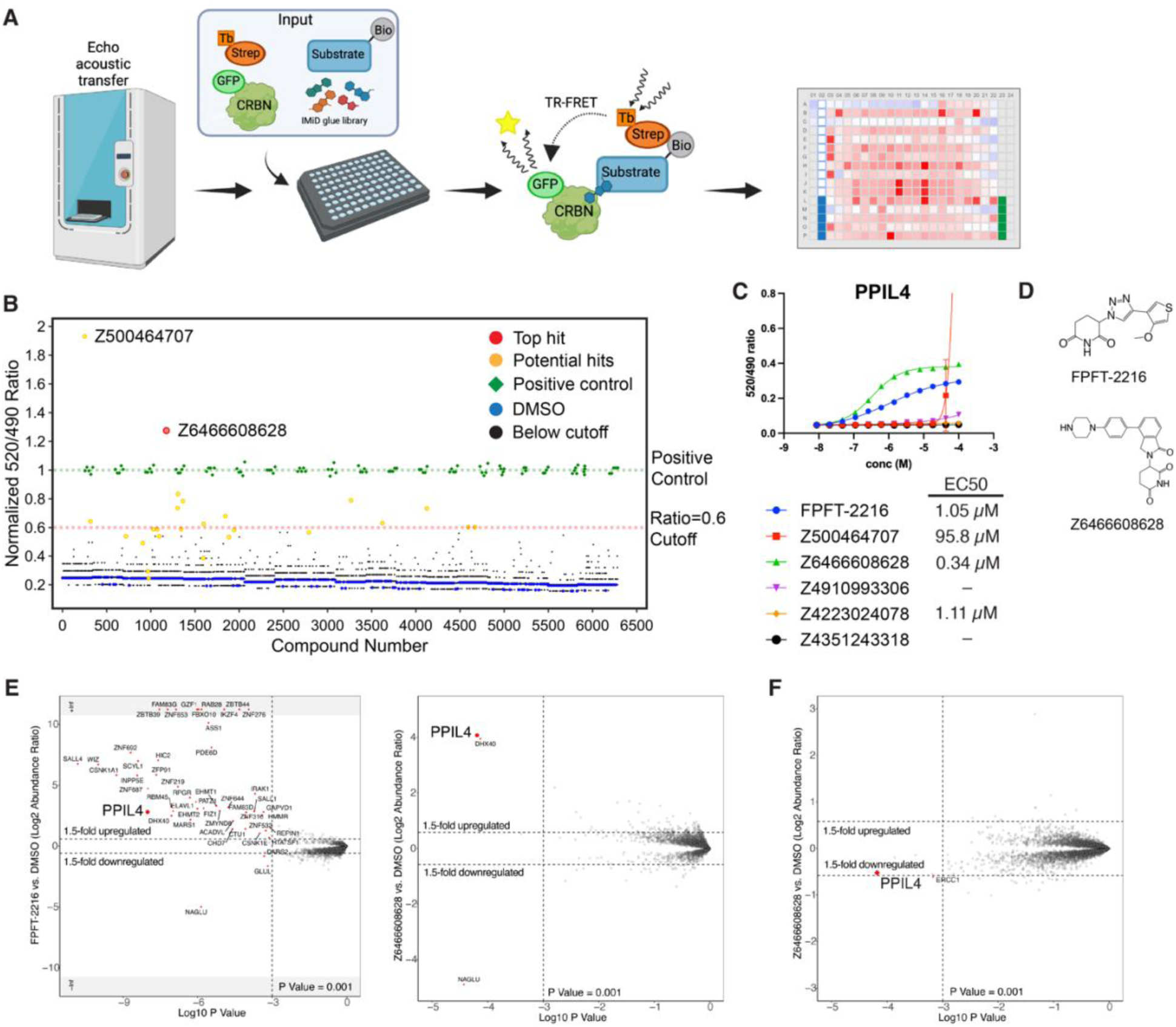
High throughput IMiD analog library screen for improved hit molecules for PPIL4. (**A**) Schematic of the high throughput TR-FRET screening workflow used to screen >6,000 IMiD analogs. (**B**) TR-FRET: Normalized 520/490 ratio for each of the >6,000 compounds derived from IMiD molecules combined from the Gray/Fischer laboratories and those purchased from Enamine with GFP-CRBN-DDB1ΔB at 50 nM, biotinylated PPIL4 at 20 nM, and terbium-streptavidin at 2 nM. (**C**) TR-FRET: titration of FPFT-2216 and lead compounds to GFP-CRBN-DDB1ΔB at (200 nM), biotinylated PPIL4 at 20 nM, and terbium-streptavidin at 2 nM. Values were determined by technical replicates of n=2. (**D**) Chemical structures of FPFT-2216 alongside new lead compound from **B** and **C**. (**E**) Scatterplot depicting relative protein abundance following Flag-CRBN-DDB1ΔB enrichment from Kelly cell in-lysate treatment with FPFT-2216 (left) and Z6466608626 (right) and recombinant Flag-CRBN-DDB1ΔB spike in. Scatterplot displays fold change in abundance to DMSO. Significant changes were assessed by moderated t-test as implemented in the limma package^80^ with log_2_ FC shown on the y-axis and negative log_10_ P-value on the x-axis. (**F**) Scatterplot depicting relative protein abundance following Z6466608626 treatment in MOLT4 cells. Significant changes were assessed by moderated t-test as implemented in the limma package^80^ with log_2_ FC shown on the y-axis and negative log_10_ P-value on the x-axis.

To test the efficacy and selectivity of our lead compound, we first performed IP-MS in comparison with FPFT-2216 in Kelly cell lysate. While FPFT-2216 recruited many proteins, Z6466608628 selectively recruited PPIL4, along with its binding partner DHX40 (**Figure 5E**, **Table S5**). We then performed global proteomics in MOLT4 cells to confirm that Z6466608628 can induce selective downregulation of PPIL4 (**Figure 5F**, **Table S5**). These data collectively demonstrate the complete workflow, starting from the identification of a novel non-ZF target PPIL4 in a chemoproteomics screen, to the discovery of a new PPIL4 selective molecular glue that would serve as an excellent lead molecule for structural optimization.

## DISCUSSION

Targeted protein degradation and induced proximity are part of a rapidly expanding field focused on the development of small molecules that leverage induced neo-protein-protein interactions to drive pharmacology. In this study, we develop and showcase a new workflow for high sensitivity, unbiased target identification of degraders and non-degrading molecular glues, identifying more than 290 targets recruited to CRBN by IMiD-like molecules. We demonstrate that this new approach to target identification can reveal critical insights and new targets that are missed by traditional screening methodologies and provide a blueprint from discovery to optimization and structure guided design of new molecular glue degraders.

Thalidomide and its derivatives, lenalidomide and pomalidomide (IMiDs), have had a checkered past. These molecules have been in use for a variety of indications, on and off, since the 1950’s and have experienced perhaps the greatest turnaround in drug history. From devastating birth defects to effective hematological cancer therapy, and more recently, significant investment in utilizing these molecules for TPD-based therapeutics. While a decade of research has slowly uncovered around 50 reported neo-substrates of IMiD’s, thousands of proteins harbor G-loops that have the potential for recruitment to CRBN by IMiD molecules. Our simple, cost effective and highly scalable unbiased screening workflow combines whole cell lysate with recombinant Flag-CRBN and degraders to enrich target binders from the complex proteome. Through an IMiD-analog diversity screen across two cell lines, we mapped a significant expansion of the neo-substrate repertoire by identifying 298 proteins recruited to CRBN, with 270 of these being novel targets. Unlike many current high throughput screening workflows that focus on the end point – degradation, this workflow allows us to explore the fundamental first step of proximity induced degradation – recruitment, where we are now able to identify targets that are directly or indirectly recruited to CRBN. This sensitive workflow sheds light onto a previously unchartered element of the molecular glue mechanism of action and establishes insights into how and why certain molecular glues may exhibit higher efficacy than others. Surprisingly, we discovered targets recruited to CRBN that are resistant to degradation, demonstrating the first examples of targets being glued to CRBN without productive degradation. Exploration of two of these targets, ASS1 and ZBED3, does not offer any clues as to why they are not degraded since both have reported ubiquitylated sites^77^. Numerous possibilities exist, from these targets being tightly preoccupied by other binding partners, geometric constraints leading to inaccessibility of lysines, removal of ubiquitylation by deubiquitinases, or preclusion of the catalytic sites due to size and shape preventing active ubiquitylation. It is important to note that the non-degrading functions of these molecular glues may have interesting degradation-independent pharmacology that have not yet been investigated, thus providing an opportunity for future experimental research.

The comprehensive G-loop database provided us with prefiltered insights as to whether these targets have the potential to be recruited to CRBN through the currently established mechanism of G-loop binding. However, although most targets identified in this study do have a structural G-loop, we do have numerous instances of proteins that do not harbor a G-loop. Some of these targets do have a structurally similar hairpin motif but are lacking the ‘essential’ glycine in position six. Whereas other targets did not have this structural motif at all. These findings indicate the potential for alternative recruitment mechanisms such as proteins piggybacking on a direct G-loop carrying target. This concept of collateral (or bystander) targeting was also demonstrated in a study exploring HDAC degradability, where it was found that both HDACs, and their known complex binding partners can be degraded^31^. Alternatively, and perhaps more intriguingly, the potential for recruitment of proteins through a distinct structural motif suggesting there may be new binding mechanisms that are pending discovery. The potential capacity for IMiDs to yield interfaces favorable for recruitment of various structural motifs would considerably expand the diversity CRBN neo-substrates and broaden therapeutic applications.

Amongst the targets identified in this study, we not only discovered many new C_2_H_2_ ZF transcription factor targets but also extended targets beyond C_2_H_2_ ZF proteins, into additional classes of proteins such as those containing RNA recognition motif (RRM) domain and kinase domains, confirming that CRBN is an incredibly versatile ligase and very well suited to hijacking for TPD applications. We reveal 251 non-ZF targets, a dramatic increase in the breadth and number of proteins targeted by CRBN from the currently reported targets of less than a dozen. Direct binding data using TR-FRET on a selection of these targets validates their direct binding mechanism, and structural characterization further corroborates this binding while validating the generated G-loop alignment database as a tool to assist prioritization of targets using clash score assessment. Using the accumulative data, we selected a novel non-ZF neo-substrate, PPIL4, for additional screening to illustrate the utility of this workflow for prioritization efforts. After a biochemical ternary complex recruitment screen of around 6,000 IMiD analogs, we selected a single hit compound and used chemoproteomics to confirm selective recruitment of PPIL4 to CRBN. Genomic studies have reported that PPIL4 is essential for brain specific angiogenesis and has implications in intracranial aneurysms^78^, and is known to regulate the catalytic activation of the spliceosome^79^. Thus, this new molecular glue could be of great interest to target the splicing pathway, in relation to intracranial aneurysms, or in other contexts.

We believe our strategic workflow and comprehensive data package, along with outlining specific applications of these, provides a valuable resource for the chemical biology, drug discovery and induced proximity communities. Importantly, the workflow is neither limited to CRBN nor to TPD, but rather can be applied to any induced proximity application. We expect the enrichment workflow will provide a blueprint for expansion into target identification for induced proximity platforms as well as further expansion of targets for protein degraders beyond molecular glues. Through initial scouting efforts on heterobifunctional degraders and additional ligases we are confident there are many novel discoveries to be made with already existing chemistry and we envision this as an evolving resource where we will continue to release data as it becomes available.

## SIGNIFICANCE

Degraders and molecular glues are small molecules that can target and promote the degradation of specific proteins providing a novel approach for modulating protein function. Currently available unbiased methods to identify targets of degraders, although successful in identifying transient and/or degraded targets, are limited in sensitivity and ability to identify direct binders of these molecules, prohibiting identification of targets that have weak expression changes or are glued and not degraded. Here, we develop an automatable high throughput method for the identification of chemically-induced binders. We demonstrate the ability to comprehensively identify new targets by identifying 298 neo-substrates of CRBN, significantly expanding the repertoire of actionable targets. We then used structural and biochemical characterization alongside a computational structural alignment workflow to validate hit targets and selected one novel target, PPIL4, to perform a focused biochemical screen for the identification of a new lead molecule. CRBN is the most targeted ligase in the TPD field, with molecules FDA approved and more in clinical trials it is important that we understand the complete cellular and molecular impact of targeting this ligase. The findings presented in this study, open a new and complementary avenue for target identification and create a valuable data resource mapping a wide range of neo-substrates of the CRBN ligase. Through expansion of the range of CRBN targets, we not only enhance our knowledge of newly druggable targets and offer new avenues for therapeutic development, but we also enhance our understanding of the molecular mechanisms and cellular pathways that are influenced by existing and future IMiD molecules providing opportunities for improved drug design.

## ACKNOWLEDGEMENTS

We thank S. Dixon-Clarke, M. Hunkeler, T. Levitz, Y. Xiong, J. Che, T. Zhang and members of the Fischer and Gray labs for helpful discussions, reagents, and support. We thank the Harvard Cryo-EM center for Structural Biology for support on data collection. Financial support for this work was provided by the National Institutes of Health (R01CA214608 and R01CA262188 (both to E.S.F.). K.B. is a Damon Runyon Fellow supported by the Damon Runyon Cancer Research Foundation (DRG-2514-24). Figures 1A, 2A and 5A were created in Biorender.

## AUTHOR CONTRIBUTIONS

K.B. designed experiments, performed structural work and biochemical assays, analyzed the data, interpreted results, wrote the manuscript. R.J.M. initiated the study, designed proteomics experiments, performed proteomics experiments, analyzed the data. S.S.R.B. performed computational alignment analysis, interpreted results. J.W.B. initiated the study and performed biochemical experiments. H.Y designed proteomics experiments. R.J.L. wrote proteomics analysis code. D.M.A performed proteomics experiments. M.L. performed immunoblots. H.Y. performed TR-FRET screen. S.O. performed TR-FRET screen. A.L.V. synthesized molecules. N.S.G. supervised experiments. K.A.D. conceived the study, designed experiments, analyzed the data, interpreted the results, wrote the manuscript and supervised the study. E.S.F. conceived the study, interpreted results, supervised and funded the study. All authors read, edited and approved the final manuscript.

## RESOURCES AVAILABILITY

### Lead Contact

Further information and requests for resources and reagents should be directed to and will be fulfilled by the Lead Contact, Eric Fischer.

### Materials Availability

Small molecules described in this study will be made available on request, upon completion of a Materials Transfer Agreement.

## EXPERIMENTAL MODEL AND STUDY PARTICIPANT DETAILS

### Mammalian Cell culture

MOLT4 and Kelly cells were cultured in RPMI-1640 media supplemented with 10% fetal bovine serum and 2 mM L-glutamine and grown in a 37 °C incubator with 5% CO_2_. MOLT4 cells are derived from 19 year old male, Kelly cells are derived from 1 year old female.

### Insect cell culture

High Five insect cells were cultured at 27 °C in SF-4 Baculo Express ICM medium (BioConcept) in suspension. Sf9 insect cells were cultured at 27 °C in ESF 921 medium (Expression Systems).

## METHOD DETAILS

### Immunoprecipitation

A total of 1 × 10^7^ cells per IP were collected and lysed in lysis buffer (50 mM Tris pH 8, 200 mM NaCl, 2 mM TCEP, 0.1% NP-40, 10 units turbonuclease/200 µL buffer, 1x cOmplete protease inhibitor tablet/5 mL buffer) and sonicated on ice for 5 rounds of 2 seconds followed by 10 second pauses at 25% amplitude. After centrifugation clarification, lysate was transferred to new lobind tubes. 20 µg of Flag-CRBN-DDB1ΔB, 1 µM of MLN4924 and CSN5i-3 (neddylation trap)^96^, and 1 µM of selected degraders or DMSO vehicle control were added to each lysate and incubated with end-over-end rotation for 1 hour in the cold room. 20 µL of pre-washed and resuspended M2-Flag magnetic bead slurry was added to each IP and incubated with end-over-end rotation for 1 hour in the cold room. Beads were washed with wash buffer (50 mM Tris pH 8, 2 mM TCEP, 0.1% NP-40, 1x cOmplete protease inhibitor tablet/5 mL buffer) containing the respective compounds followed by specific elution method for the next application (immunoblot or mass spectrometry).

Samples prepared on the OT2 followed the same procedures as above but using scaled down equivalents of reagents. After the three detergent washes described above, the samples were washed an additional three times with non-detergent buffer (50 mM Tris pH 8, 2 mM TCEP, 1x cOmplete protease inhibitor tablet/5 mL buffer) containing the required compounds prior to appropriate elution described below.

### Immunoblot

IP samples were eluted by resuspension in SDS sample buffer and heated at 95 °C for 5 minutes. Samples were run on 4-20% Mini-PROTEAN TGX Precast Protein gels (Bio-rad). Protein was transferred to PVDF membranes using the iBlot 2.0 dry blotting system (Thermo Fisher Scientific). Membranes were blocked with Intercept blocking solution (LiCor), washed three times with PBST and incubated with primary antibodies diluted using Intercept® T20 (PBS) Antibody Diluent, overnight at 4 °C, followed by three washes with PBST and incubation with secondary antibodies for 1 hour in the dark. The membrane was washed three final times in PBST and imaged on the Odyssey LCx imaging system (LiCor).

### Cloning and Protein Expression

All proteins are derived from human origin sequences. ZBED3 (residues 39-108), ZNF219 (residues 53-110) were cloned into pGEX4T1-TEV vectors with C-terminal Avi-Strep fusion. These were expressed in LOBSTR BL21(DE3) *E. coli*, purified by GST-affinity, followed by liberation of the GST-tag by overnight TEV protease incubation, ion exchange chromatography, and size exclusion chromatography into a final buffer of 25 mM HEPES, 200 mM NaCl, and 1 mM TCEP, pH 7.5. UBE2D3, UBE2G1, and UBE2M were cloned into pGEX4T1-TEV vectors, and NEDD8 was cloned into pGEX4T1-3C vector. Purification of these proteins were performed as described above. APPBP1-UBA3 was cloned into a pET based vector and expressed in LOBSTR BL21(DE3) *E. coli*, purified by nickel affinity, ion exchange chromatography, and size exclusion chromatography into a final buffer of 25 mM HEPES, 200 mM NaCl, and 1 mM TCEP, pH 7.5. Ubiquitin was cloned into a pET3a vector and was expressed in Rosetta BL21(DE3) *E. coli*. Ubiquitin was purified by SP Sepharose resin at pH 4, and was subject to size exclusion chromatography into a final buffer of 25 mM HEPES, 150 mM NaCl, 1 mM TCEP, pH 7.5. PDE6D, DTWD1, PPIL4, PPIL4 RRM (residues 240-318), and RAB28 were cloned into pAC8-Strep-Avi vectors. Baculovirus was generated in *Spodoptera frugiperda* cells, and proteins were expressed in *Trichoplusia ni* cells. These were purified by Strep-affinity, followed by ion exchange chromatography and size exclusion chromatography into 25 mM HEPES, 200 mM NaCl, and 1 mM TCEP, pH 7.5. GST-TEV-CRBN, GST-TEV-eGFP-CRBN, DDB1 FL, GST-TEV-UBA1, CUL4A (residues 38-C), RBX1(residues 5-C) were cloned into pLIB vectors. His-TEV-DDB1ΔBPB (Δresidues 396-705 with a GNGNSG linker) and Flag-Spy-CRBN were cloned into a pAC8 vector. CRBN and DDB1 were coexpressed in the following pairs, CRBN-DDB1, CRBN-DDB1ΔB, Flag-CRBN-DDB1ΔB, and eGFP-CRBN-DDB1ΔB, purified by GST-affinity, followed by TEV cleavage overnight, ion exchange chromatography, and size exclusion chromatography into 25 mM HEPES, 150 mM NaCl, 1 mM TCEP, pH 7.5. CUL4A and RBX1 were coexpressed, CUL4-RBX1, and UBA1 were purified as described above.

### Time-Resolved Fluorescence Energy Transfer (TR-FRET)

Ternary complex formation of CRBN-DDB1, neo-substrate, and compound was measured by TR-FRET. 20 nM biotinylated neo-substrate, 2 nM Tb (CoraFluor-1)-labeled Streptavidin, and 200 nM eGFP-CRBN-DDB1ΔB were added in assay buffer (25 mM HEPES, 100 mM NaCl, 0.5% Tween-20, and 0.5% BSA). This reaction mix was added to a 384 well microplate, and compound was titrated to indicated concentrations using a D300e Digital Dispenser (HP). Reaction was incubated at room temperature for 1 hour and was measured using a PHERAstar FS microplate reader (BMG Labtech). The 520 nm/490 nm signal ratio was measured to calculate ternary complex formation, and datapoints were plotted to calculate EC50 values by using agonist versus response variable slope (four parameter model) in Graphpad Prism (v10.1.1).

### In-vitro ubiquitylation assay

Ubiquitylation of neo-substrates was performed by premixing 500 nM neddylated CRL4^CRBN^, 2 µM UBE2D3, 2 µM UBE2G1, 200 nM UBA1, 500 nM Strep tagged-neo-substrate, and 10 µM compound in assay buffer (25 mM HEPES, 100 mM NaCl, 10 mM MgCl_2_, 5 mM ATP, pH 7.5). This mixture was incubated for 15 min on ice, and reaction was started by adding 60 µM ubiquitin at room temperature. Sample was quenched by addition of SDS sample buffer and reaction products were separated on 4-20% SDS-PAGE gels. Assay was analyzed by immunoblotting as described above. Neddylation of CUL4-RBX1 was performed as described in Scott et al., 2014^97^. In brief, 12 µM CUL4-RBX1, 1 µM UBE2M, 0.2 µM APPBP1-UBA3, 25 µM NEDD8 was incubated at room temperature for 10 min in assay buffer (25 mM HEPES, 100 mM NaCl, 10 mM MgCl_2_, 5 mM ATP, pH 7.5). Reaction was quenched by addition of 10mM DTT and was purified by size exclusion chromatography into buffer containing 25 mM HEPES, 150 mM NaCl, 1 mM TCEP, pH 7.5.

### Cryo-EM sample preparation and Data processing

10 µM CRBN-DDB1ΔB, 20 µM PPIL4 RRM, and 100 µM FPFT-2216 were incubated on ice for 30 min in buffer containing 25 mM HEPES, 150 mM NaCl, 1 mM TCEP, pH 7.5. The protein complex was diluted to 1.5 µM final concentration before application to grids. Quantifoil UltraAuFoil grids (R0.6/1) were glow discharged at 20 mA for 2 min. Prior to complex application, 4 µL of 10 µM CRBN-agnostic IKZF1 (residues 140-196, harboring mutants Q146A, G151N) was applied to saturate the air-water interface^82^. After 1 min, the grid was blotted from the back, and complex was applied, and immediately plunged into liquid ethane.

Dataset was collected on a Talos Arctica at 200 kV equipped with a Gatan K3 direct detector in counting mode. Movies were collected with a total dose of 54 e^-^/Å^2^ over 40 frames, at 1.1 Å/pixel with a nominal magnification of 36,000x, with a defocus range of -0.8 µm to -2.0 µm.

### Sample preparation for immunoprecipitation mass spectrometry (IP-MS)

Immunoprecipitation (IP) was performed as described above. After the final wash step, samples were eluted using 0.1 M Glycine-HCl, pH 2.7. Tris (1M, pH 8.5) was added to elution to reach a pH of 8. Samples were then reduced with 10 mM TCEP for 30 min at room temperature, followed by alkylation with 15 mM iodoacetamide for 45 min at room temperature in the dark. Alkylation was quenched by the addition of 10 mM DTT. Proteins were isolated by methanol-chloroform precipitation (only for manual IPs. Automated IPs undergo three non-detergent washes instead). The protein pellets were dried and then resuspended in 50 μL 200 mM EPPS pH 8.0. The resuspended protein samples were digested with 2 μg LysC and 1 μg Trypsin overnight at 37°C. Sample digests were acidified with formic acid to a pH of 2-3 prior to desalting using C18 solid phase extraction plates (SOLA, Thermo Fisher Scientific). Desalted peptides were dried in a vacuum-centrifuged and reconstituted in 0.1% formic acid for LC-MS analysis.

### Label free quantitative mass spectrometry with DDA and data analysis

Data were collected using an Orbitrap Exploris 480 mass spectrometer (Thermo Fisher Scientific) equipped with a FAIMS Pro Interface and coupled with a UltiMate 3000 RSLCnano System. Peptides were separated on an Aurora 25 cm x 75 μm inner diameter microcapillary column (IonOpticks), and using a 60 min gradient of 5 -25% acetonitrile in 1.0% formic acid with a flow rate of 250 nL/min. Each analysis used a TopN data-dependent method. The data were acquired using a mass range of m/z 350 – 1200, resolution 60,000, 300% normalized AGC target, auto maximum injection time, dynamic exclusion of 30 sec, and charge states of 2-6. TopN 40 data-dependent MS2 spectra were acquired with a scan range starting at m/z 110, resolution 15,000, isolation window of 1.4 m/z, normalized collision energy (NCE) set at 30%, standard AGC target and the automatic maximum injection time.

Proteome Discoverer 2.5 (Thermo Fisher Scientific) was used for .RAW file processing and controlling peptide and protein level false discovery rates, assembling proteins from peptides, and protein quantification from peptides. MS/MS spectra were searched against a Swissprot human database (January 2021) with both the forward and reverse sequences as well as known contaminants such as human keratins. Database search criteria were as follows: tryptic with two missed cleavages, a precursor mass tolerance of 10 ppm, fragment ion mass tolerance of 0.03 Da, static alkylation of cysteine (57.0215 Da) and variable oxidation of methionine (15.995 Da), N-terminal acetylation (42.011 Da) and phosphorylation of serine, threonine and tyrosine (75.966 Da). Peptides were quantified using the MS1 Intensity, and peptide abundance values were summed to yield the protein abundance values. Resulting data was filtered to only include proteins that had a minimum of 3 counts in at least 3 replicates of each independent comparison of treatment sample to the DMSO control. Protein abundances were globally normalized and scaled using in-house scripts in the R framework (R Development Core Team, 2014). Proteins with missing values were imputed by random selection from a Gaussian distribution either with a mean of the non-missing values for that treatment group or with a mean equal to the median of the background (in cases when all values for a treatment group are missing). Significant changes comparing the relative protein abundance of these treatment to DMSO control comparisons were assessed by moderated t-test as implemented in the limma package within the R framework^80^.

### Label free quantitative mass spectrometry with diaPASEF and data analysis

Data were collected using a TimsTOF Pro2 (Bruker Daltonics, Bremen, Germany) coupled to a nanoElute LC pump (Bruker Daltonics, Bremen, Germany) via a CaptiveSpray nano-electrospray source. Peptides were separated on a reversed-phase C18 column (25 cm x 75 μM ID, 1.6 μM, IonOpticks, Australia) containing an integrated captive spray emitter. Peptides were separated using a 50 min gradient of 2 - 30% buffer B (acetonitrile in 0.1% formic acid) with a flow rate of 250 nL/min and column temperature maintained at 50 °C. To perform diaPASEF, the precursor distribution in the DDA m/z-ion mobility plane was used to design an acquisition scheme for DIA data collection which included two windows in each 50 ms diaPASEF scan. Data was acquired using sixteen of these 25 Da precursor double window scans (creating 32 windows) which covered the diagonal scan line for doubly and triply charged precursors, with singly charged precursors able to be excluded by their position in the m/z-ion mobility plane. These precursor isolation windows were defined between 400 - 1200 m/z and 1/k0 of 0.7 - 1.3 V.s/cm2.

The diaPASEF raw file processing and controlling peptide and protein level false discovery rates, assembling proteins from peptides, and protein quantification from peptides was performed using library free analysis in DIA-NN 1.8^98^. Library free mode performs an in-silico digestion of a given protein sequence database alongside deep learning-based predictions to extract the DIA precursor data into a collection of MS2 spectra. The search results are then used to generate a spectral library which is then employed for the targeted analysis of the DIA data searched against a Swissprot human database (January 2021). Database search criteria largely followed the default settings for directDIA including: tryptic with two missed cleavages, carbamidomethylation of cysteine, and oxidation of methionine and precursor Q-value (FDR) cut-off of 0.01. Precursor quantification strategy was set to Robust LC (high accuracy) with RT-dependent cross run normalization. Resulting data was filtered to only include proteins that had a minimum of 3 counts in at least 4 replicates of each independent comparison of treatment sample to the DMSO control. Protein abundances were globally normalized using in-house scripts in the R framework (R Development Core Team, 2014). Proteins with missing values were imputed by random selection from a Gaussian distribution either with a mean of the non-missing values for that treatment group or with a mean equal to the median of the background (in cases when all values for a treatment group are missing). Significant changes comparing the relative protein abundance of these treatment to DMSO control comparisons were assessed by moderated t-test as implemented in the limma package within the R framework^80^.

### TMT quantitative global proteomics

Cells were lysed by the addition of lysis buffer (8 M urea, 50 mM NaCl, 50 mM 4-(2-hydroxyethyl)-1-piperazineethanesulfonic acid (EPPS) pH 8.5, Protease and Phosphatase inhibitors) followed by manual homogenization by 20 passes through a 21-gauge (1.25 in. long) needle. Lysate was clarified by centrifugation and protein quantified using bradford (Bio-Rad) assay. 100 µg of protein for each sample was reduced, alkylated and precipitated using methanol/chloroform as previously described^19^ and the resulting washed precipitated protein was allowed to air dry. Precipitated protein was resuspended in 4 M urea, 50 mM HEPES pH 7.4, buffer for solubilization, followed by dilution to 1 M urea with the addition of 200 mM EPPS, pH 8. Proteins were digested for 12 hours at room temperature with LysC (1:50 ratio), followed by dilution to 0.5 M urea and a second digestion step was performed by addition of trypsin (1:50 ratio) for 6 hours at 37 °C. Anhydrous ACN was added to each peptide sample to a final concentration of 30%, followed by addition of Tandem mass tag (TMT) reagents at a labelling ratio of 1:4 peptide:TMT label. TMT labelling occurred over a 1.5 hour incubation at room temperature followed by quenching with the addition of hydroxylamine to a final concentration of 0.3%. Each of the samples were combined using adjusted volumes and dried down in a speed vacuum followed by desalting with C18 SPE (Sep-Pak, Waters). The sample was offline fractionated into 96 fractions by high pH reverse-phase HPLC (Agilent LC1260) through an aeris peptide xb-c18 column (phenomenex) with mobile phase A containing 5% acetonitrile and 10 mM NH_4_HCO_3_ in LC-MS grade H_2_O, and mobile phase B containing 90% acetonitrile and 5 mM NH4HCO3 in LC-MS grade H_2_O (both pH 8.0). The resulting 96 fractions were recombined in a non-contiguous manner into 24 fractions and desalted using C18 solid phase extraction plates (SOLA, Thermo Fisher Scientific) followed by subsequent mass spectrometry analysis.

Data were collected using an Orbitrap Fusion Lumos mass spectrometer (Thermo Fisher Scientific, San Jose, CA, USA) coupled with a Proxeon EASY-nLC 1200 LC system (Thermo Fisher Scientific, San Jose, CA, USA). Peptides were separated on a 1) 50 cm 75 μm inner diameter EasySpray ES903 microcapillary column (Thermo Fisher Scientific). Peptides were separated over a 190 min gradient of 6 -27% acetonitrile in 1.0% formic acid with a flow rate of 300 nL/min. Quantification was performed using a MS3-based TMT method as described previously^99^. The data were acquired using a mass range of m/z 340 – 1350, resolution 120,000, AGC target 5 x 105, maximum injection time 100 ms, dynamic exclusion of 120 seconds for the peptide measurements in the Orbitrap. Data dependent MS2 spectra were acquired in the ion trap with a normalized collision energy (NCE) set at 35%, AGC target set to 1.8 x 104 and a maximum injection time of 120 ms. MS3 scans were acquired in the Orbitrap with HCD collision energy set to 55%, AGC target set to 2 x 105, maximum injection time of 150 ms, resolution at 50,000 and with a maximum synchronous precursor selection (SPS) precursors set to 10.

### TMT quantitative global proteomics LC-MS data analysis

Proteome Discoverer 2.2 (Thermo Fisher Scientific) was used for .RAW file processing and controlling peptide and protein level false discovery rates, assembling proteins from peptides, and protein quantification from peptides. The MS/MS spectra were searched against a Swissprot human database (January 2021) containing both the forward and reverse sequences. Searches were performed using a 20 ppm precursor mass tolerance, 0.6 Da fragment ion mass tolerance, tryptic peptides containing a maximum of two missed cleavages, static alkylation of cysteine (57.02146 Da), static TMT labelling of lysine residues and N-termini of peptides (229.1629 Da), and variable oxidation of methionine (15.99491 Da). TMT reporter ion intensities were measured using a 0.003 Da window around the theoretical m/z for each reporter ion in the MS3 scan. The peptide spectral matches with poor quality MS3 spectra were excluded from quantitation (summed signal-to-noise across channels < 110 and precursor isolation specificity < 0.5), and the resulting data was filtered to only include proteins with a minimum of 2 unique peptides quantified. Reporter ion intensities were normalized and scaled using in-house scripts in the R framework^100^. Statistical analysis was carried out using the limma package within the R framework^80^.

### Searching for CRBN-compatible G-loops in the AlphaFold2 database

Using MASTER v1.6^61^, the AlphaFold2 human database (v4)^62^ comprising 23,391 protein structures was queried for 8-residue loops with backbone root-mean-squared-deviation less than 1.5 Å to CK1α residues 35-42 (extracted from PDB 5FQD^12^. Loops not having a glycine at the sixth residue were filtered out resulting in a set of 46,040 G-loops from 10,926 proteins. Next, domains containing the G-loops were extracted using domain definitions from DPAM^63^ and aligned to the casein kinase I α reference loop and CRBN from 5FQD. To estimate backbone clashes, each structure was coarse-grained to represent the side chain as a pseudoatom and scored using Rosetta v13.3^64^. Coarse-graining the structure makes the score rotamer-independent. The interchain_vdw term of the score function was used as a clash score and represents a modified Lennard-Jones 6–12 potential that penalizes atoms overlapping at the interface^101^. Domains with a clash score of greater than 200 were filtered out resulting in a list 16,0901 loops from 7,111 proteins.

### Relieving minor clashes with Rosetta refinement

For select domains with low clash scores, clashes were relieved by relaxing the neo-substrate with Rosetta FastRelax^64,65^ while holding the G-loop in place. No energy minimization or rotamer optimization was performed on CRBN. Rigid body translation between CRBN and the neo-substrate was disabled. Of the 10 independent trajectories run, the one with the lowest total score was selected and then the clash score was calculated as above.

### Quantification and statistical analysis

Statistical methods are described in the according figure legends, and methods. Cryo-EM statistics are based on the gold-standard FSC=0.143 to determine resolution. For quantitative proteomics experiments, statistical analysis was carried out using the limma package within the R framework^80^.

---

**Figure S1.**
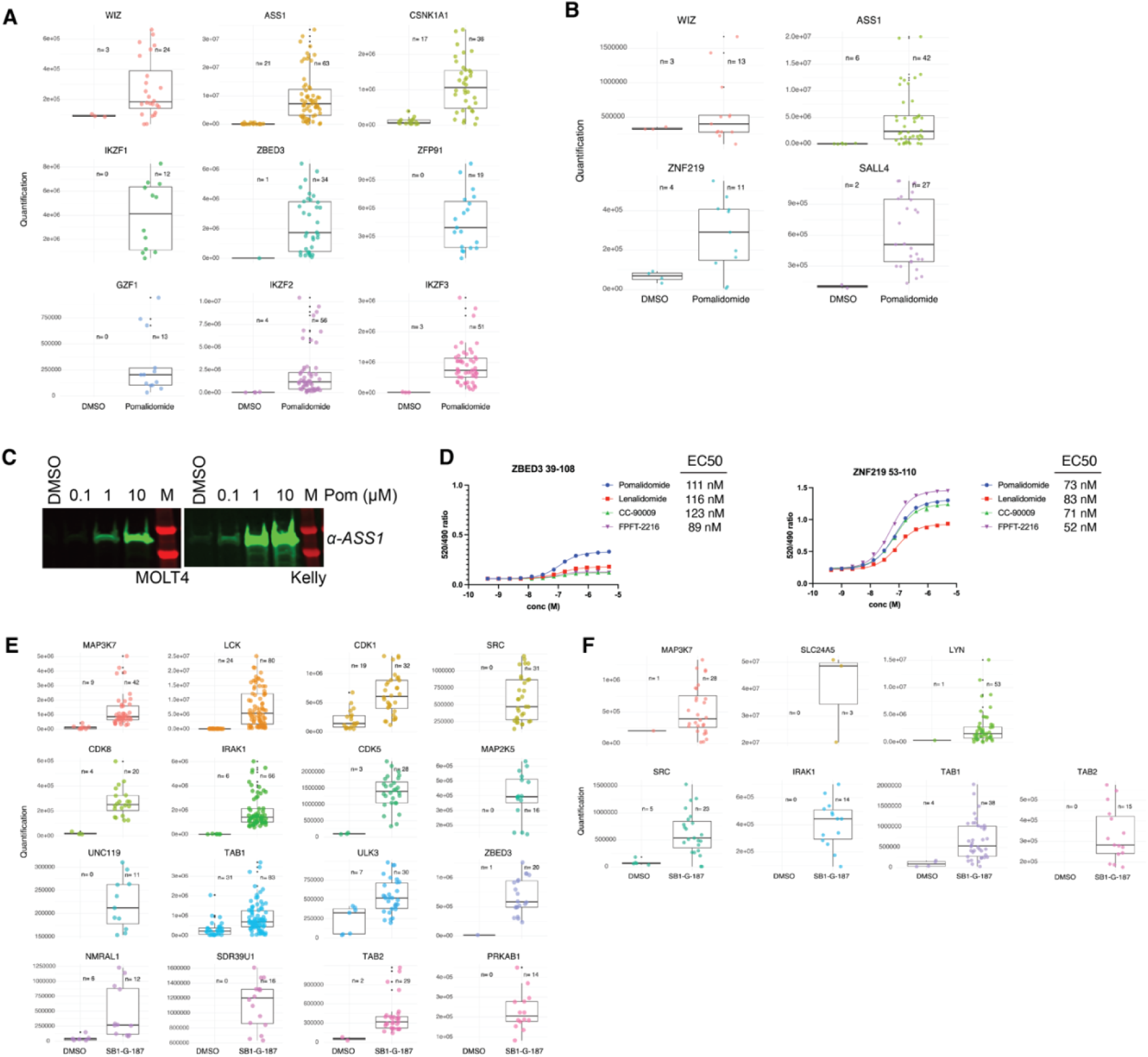
Validation of enriched targets from lysate IPs, related to Figure 1, Table S1. (**A**) Box plot depicting all quantified peptides for each of the enriched targets from MOLT4 cells comparing DMSO control and Pomalidomide treatments. (**B**) As in **A** but for Kelly cells. (**C**) Flag-CRBN IP experiments were performed in the presence of increasing concentration of pomalidomide in both MOLT4 and Kelly cells. Following elution, ASS1 protein levels were assessed by immunoblot. (**D**) TR-FRET: titration of IMiD analogs to GFP-CRBN-DDB1ΔB at 200 nM, ZBED3_39-108_ or ZNF219_53-110_ at 20 nM, and terbium-streptavidin at 2 nM. Values were determined by technical replicates of n=2. (**E**) Box plot depicting all quantified peptides for each of the enriched targets from MOLT4 cells comparing DMSO control and SB1-G-187 treatments. (**F**) As in **E** but for Kelly cells.

**Figure S2.**
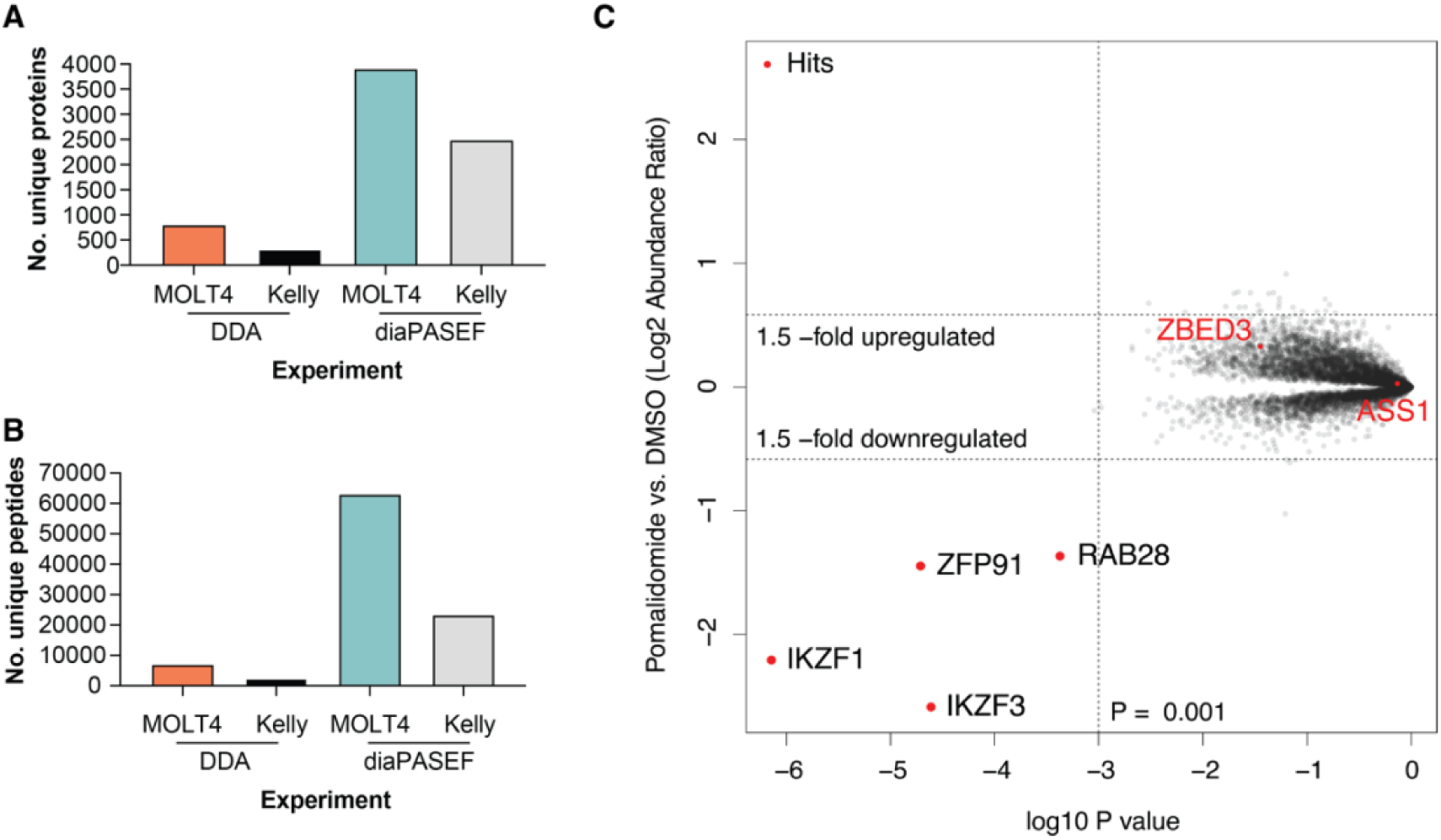
Quantitative proteomics for exploration of targets recruited by IMiD analogs, related to Figure 2. (**A**) Histogram displaying number of unique proteins quantified in DDA and diaPASEF IP-MS experiments for MOLT4 and Kelly cells. (**B**) Histogram displaying number of unique peptides quantified in DDA and diaPASEF IP-MS experiments for MOLT4 and Kelly cells. (**C**) Scatterplot depicting relative protein abundance following 5 µM Pomalidomide treatment in MOLT4 cells. Significant changes were assessed by moderated t-test as implemented in the limma package^80^ with log_2_ FC shown on the y-axis and negative log_10_ P-value on the x-axis.

**Figure S3.**
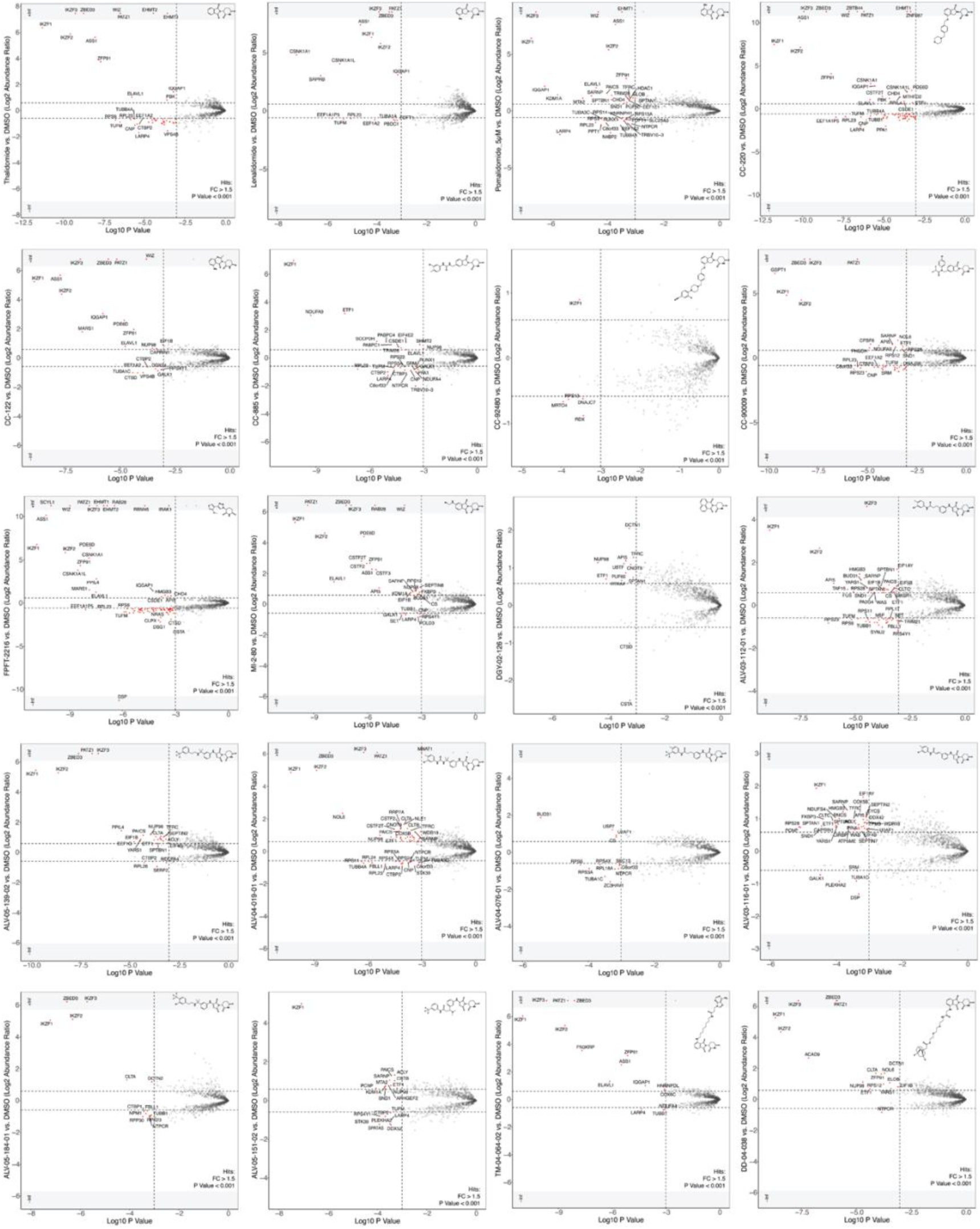
Scatterplots of IP-MS with molecular glues, related to Figure 2, 3, S4, Table S2, S3, S4. Scatterplots depicting relative protein abundance following Flag-CRBN-DDB1 enrichment from in-lysate treatment with degrader and recombinant Flag-CRBN-DDB1 spike in. Scatterplot displays fold change in abundance for each of the 20 molecules relative to DMSO in MOLT4 cells. Significant changes were assessed by moderated t-test as implemented in the limma package^80^ with log_2_ FC shown on the y-axis and negative log_10_ P-value on the x-axis.

**Figure S4.**
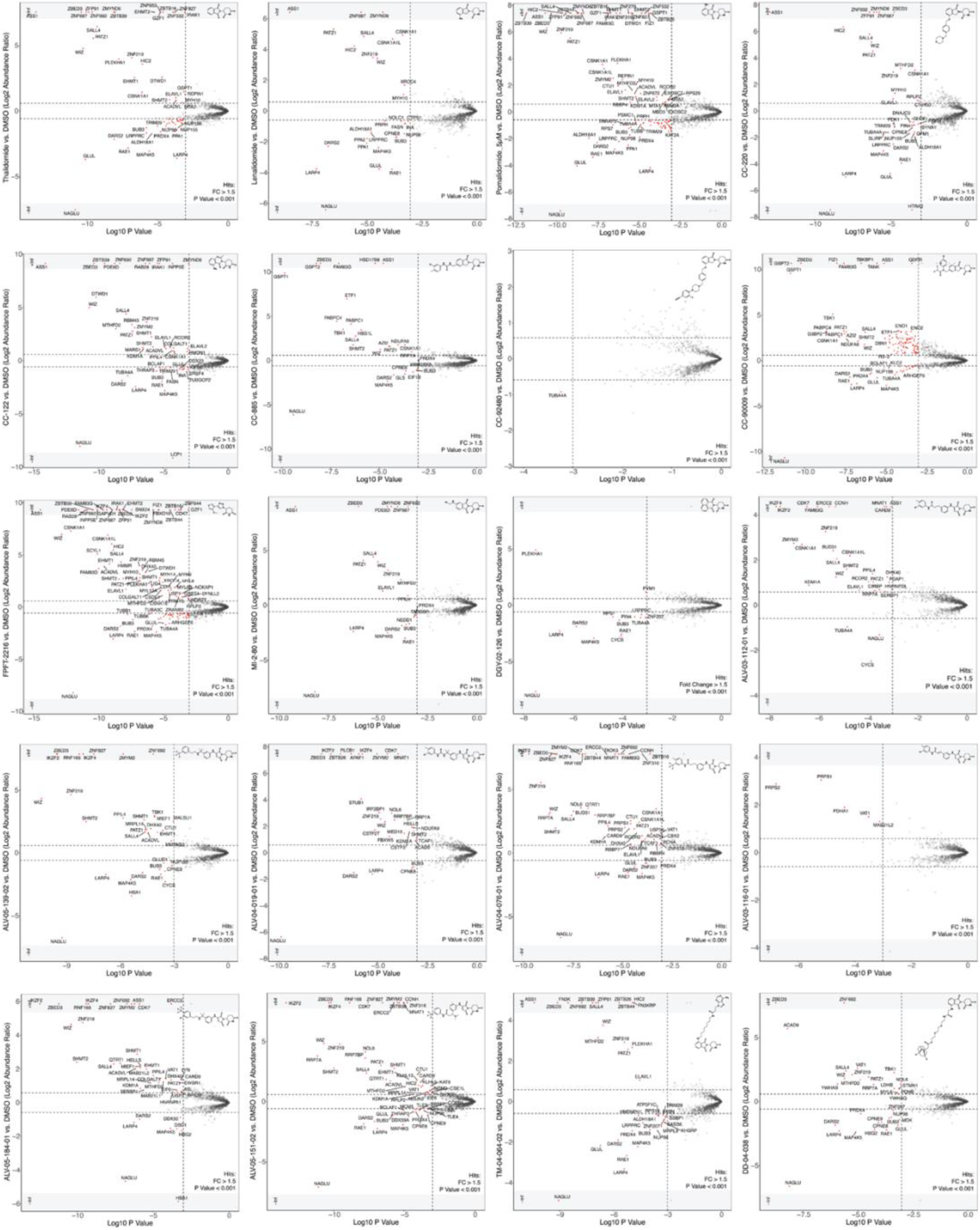
Scatterplots of IP-MS with molecular glues, related to Figure 2, 3, S4, Table S2, S3, S4. Scatterplots depicting relative protein abundance following Flag-CRBN-DDB1 enrichment from in-lysate treatment with degrader and recombinant Flag-CRBN-DDB1 spike in. Scatterplot displays fold change in abundance for each of the 20 molecules relative to DMSO in Kelly cells. Significant changes were assessed by moderated t-test as implemented in the limma package^80^ with log_2_ FC shown on the y-axis and negative log10 P-value on the x-axis.

**Figure S5.**
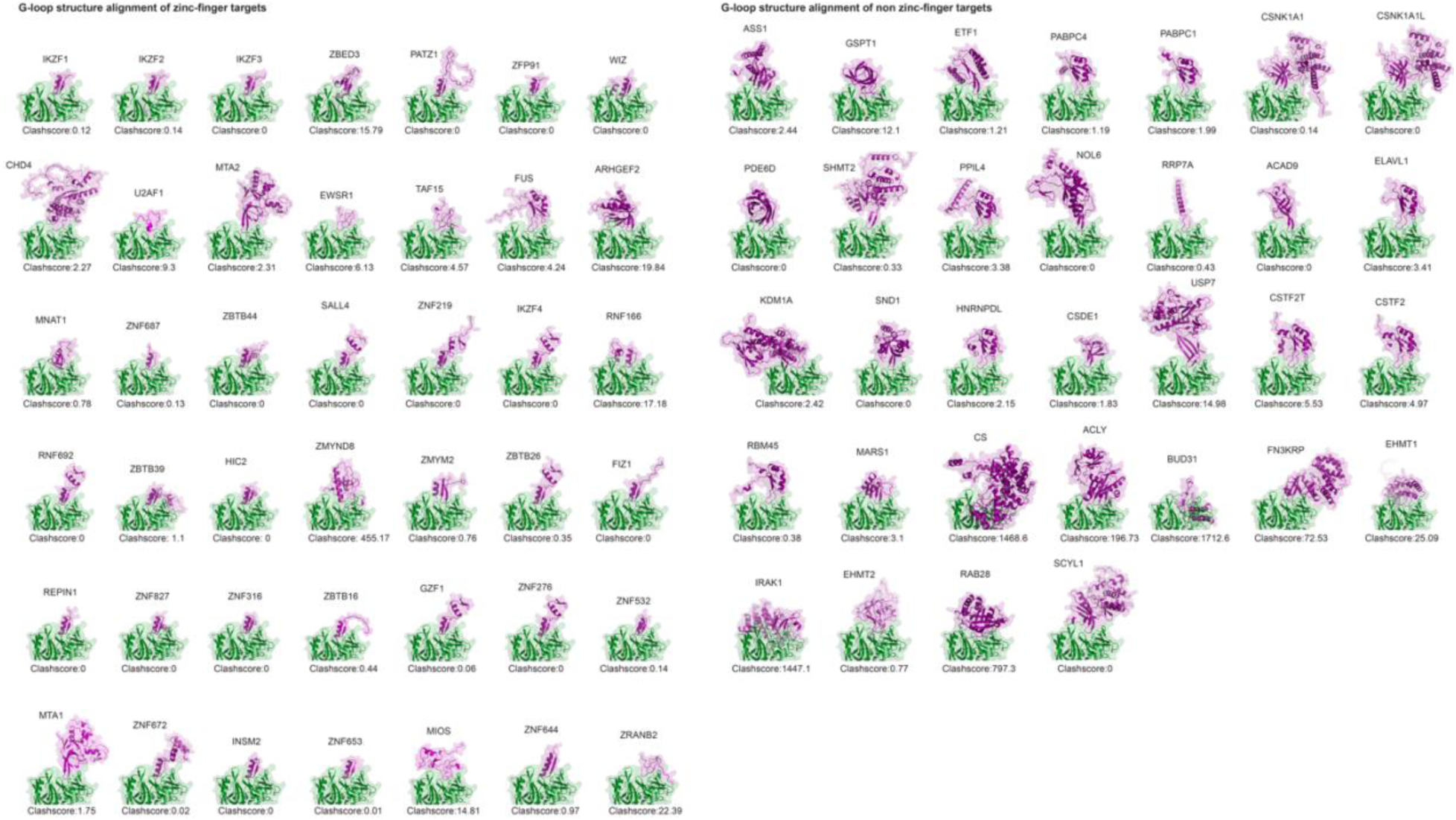
Structural alignments of G-loop neo-substrates, related to Figure 3, 4, S5, Table S5. Structural G-loop alignment showing alignment of AF2 structures by the G-loop for a subset of the neo-substrate targets identified as hits in this study with CRBN (PDB ID: 5FQD). Corresponding clash scores are displayed.

**Figure S6.**
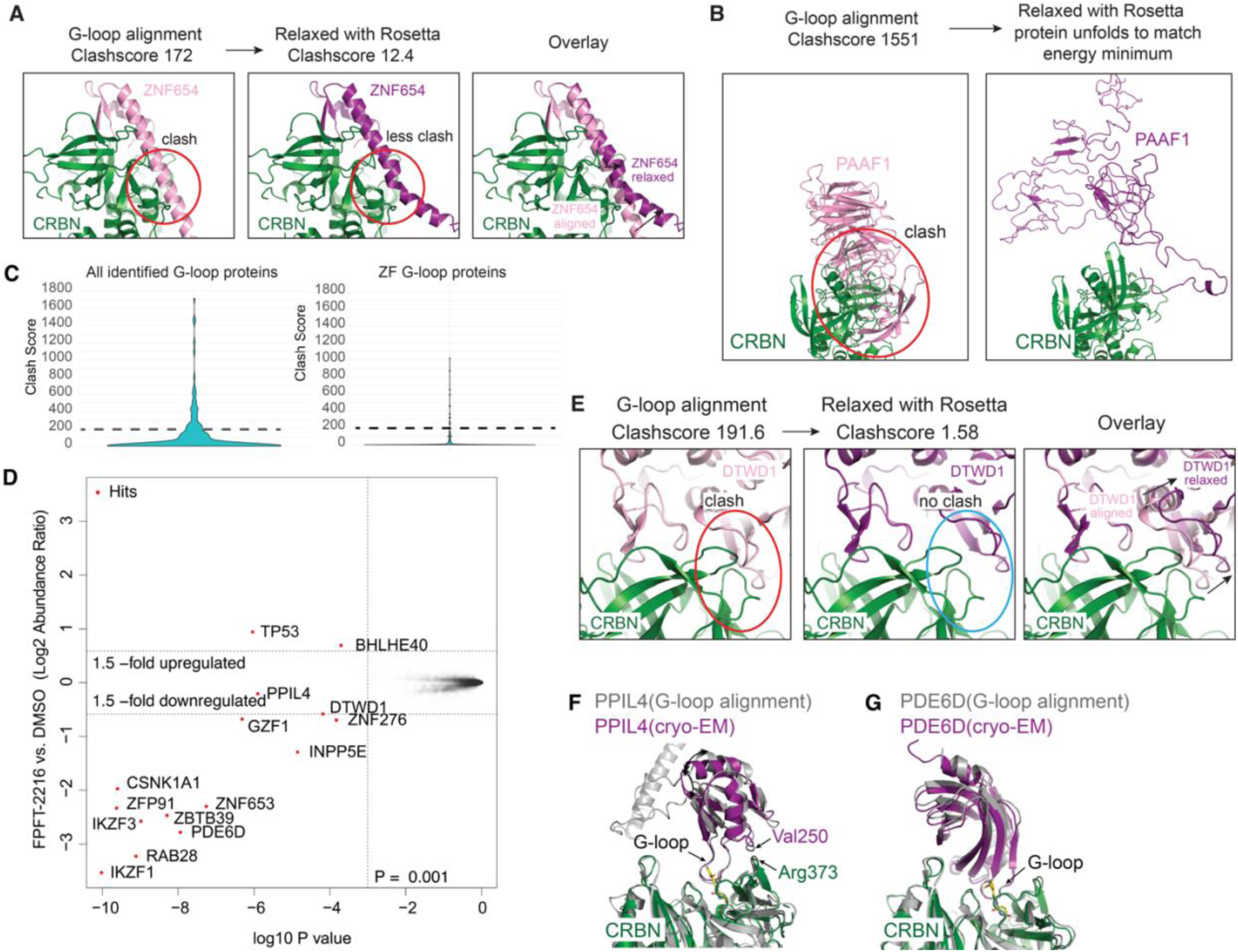
Resolving clashes from G-loop alignments, related to Figure 3, 4, S5, Table S5. (**A**) G-loop aligned AF2 structures for ZNF654 demonstrating the clash score and structural shift before and after relaxation with Rosetta. (**B**) As in **A**, but for PAAF1. (**C**) Violin plot depicting the distribution of class scores for a single G-loop from each of the identified targets in this dataset (left), and those for ZF proteins only (right). (**D**) Scatterplot depicting relative protein abundance following FPFT-2216 treatment in MOLT4 cells. Significant changes were assessed by moderated t-test as implemented in the limma package^80^ with log_2_ FC shown on the y-axis and negative log_10_ P-value on the x-axis. (**E**) As in **A**, but for DTWD1. (**F**) Overlay of G-loop aligned AF2 structure of PPIL4 and structure obtained by cryo-EM. (**G**) Same as in (**F**) but with PDE6D.

**Figure S7.**
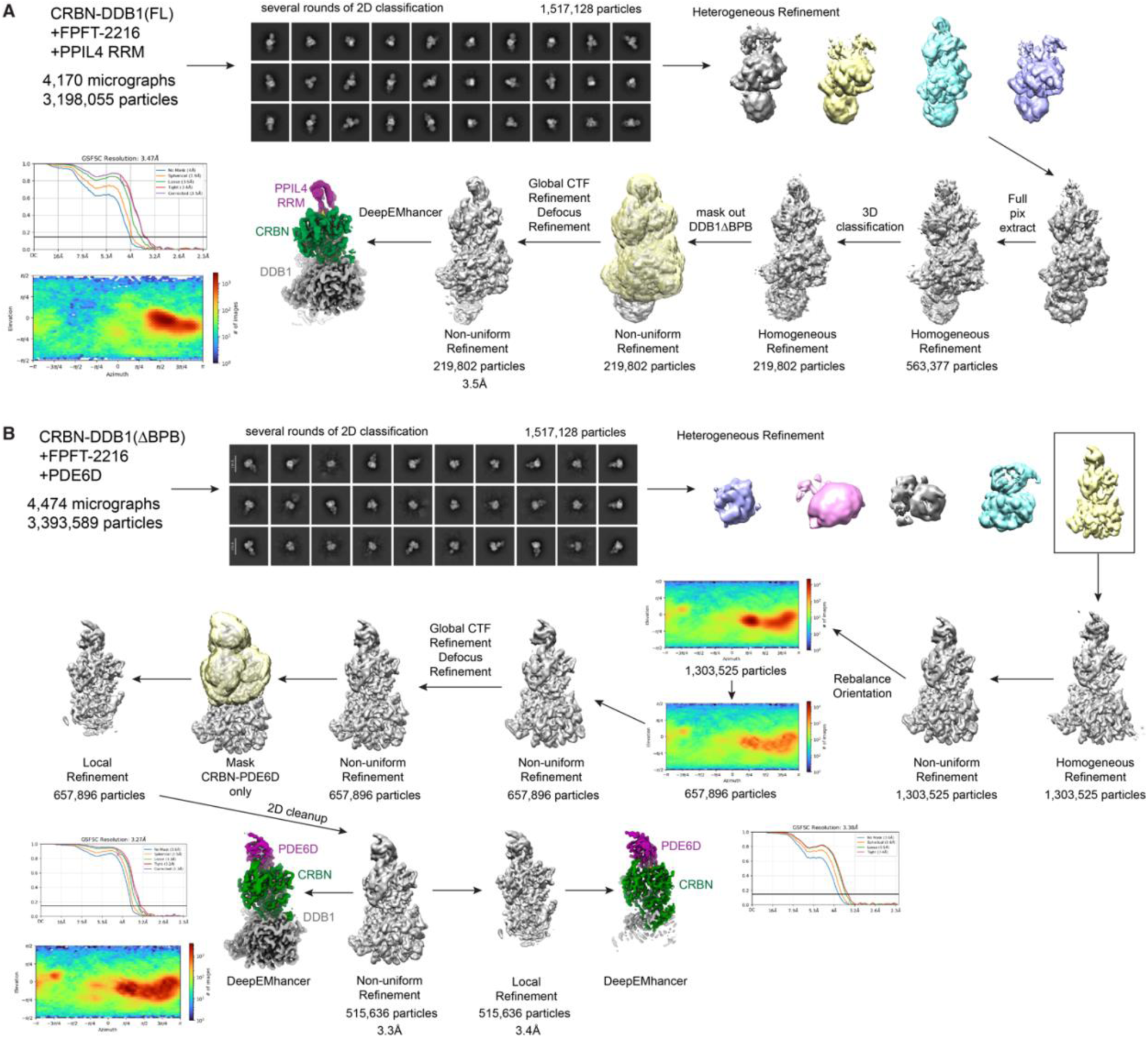
Cryo-EM processing schematic of neosubstrate-molecular glue-CRBN-DDB1, related to Figure 4, Table 1. (**A**) Cryo-EM processing schematic of CRBN-DDB1 (FL) with PPIL4 RRM domain and FPFT-2216. **(B)** Cryo-EM processing schematic of CRBN-DDB1ΔB with PDE6D and FPFT-2216.

